# CLEM*Site*, a software for automated phenotypic screens using light microscopy and FIB-SEM

**DOI:** 10.1101/2021.03.19.436113

**Authors:** José M. Serra Lleti, Anna M. Steyer, Nicole L. Schieber, Beate Neumann, Christian Tischer, Volker Hilsenstein, Mike Holtstrom, David Unrau, Robert Kirmse, John M. Lucocq, Rainer Pepperkok, Yannick Schwab

**Author notes:** contributed equally (José M. Serra Lleti, Anna M. Steyer, Nicole L. Schieber).

## Abstract

Correlative light and electron microscopy (CLEM) combines two imaging modalities, balancing out the limits of one technique with the other. In recent years, Focused Ion Beam Scanning Electron Microscopy (FIB-SEM) has emerged as a flexible method that enables semi-automated volume acquisition at the ultrastructural level. We present a toolset for adherent cultured cells that enables tracking and finding cell regions previously identified in light microscopy, in the FIB-SEM along with automatic acquisition of high-resolution volume datasets. We detect a grid pattern in both modalities (LM and EM), which identifies common reference points. The novel combination of these techniques enables complete automation of the workflow. This includes setting the coincidence point of both ion and electron beams, automated evaluation of the image quality and constantly tracking the sample position with the microscope’s field of view reducing or even eliminating operator supervision. We show the ability to target the regions of interest in EM within 5 *µ*m accuracy, while iterating between different targets and implementing unattended data acquisition. Our results demonstrate that executing high throughput volume acquisition in electron microscopy is possible.

## Introduction

Electron microscopy (EM) of cultured cells provides unique access to detailed subcellular architectures at nanometer scale. Sampling strategies are essential to ensure an accurate morphometric evaluation of subcellular phenotypes. In cases where cells are homogeneous, random sampling guarantees optimal selection of the overall population ^1,2,3^. However, different paradigms are necessary to measure sub-cellular morphologies on heterogeneous cell cultures ^4,5^. Increasing imaging throughput is one way to address heterogeneity but EM rarely achieves sufficient regimes. Correlative light and electron microscopy (CLEM) is an efficient solution to overcome such heterogeneity in EM. It capitalizes on the power of light microscopy (LM) to screen large samples for choosing cell sub-populations of interest. By applying a selection process on the light microscopy level, analysis can be focused towards specific individual cells, even if the phenotype of interest is extremely rare. Thus, various targeting strategies have been developed since the very first CLEM was performed on cultured cells ^6,7^. Individual areas of interest inside the sample can be tagged by means of laser etched frames ^8^ or cells can be seeded onto dedicated substrates that incorporate a coordinate system ^9,10^. In both cases, object correlation is established using landmarks created with artificial fiducial markers that are easily identifiable in both LM and EM. Over the years, various solutions have been developed to imprint such fiducials, such as gold or ink printing ^11,12^, laser or scalpel etching ^9,13^ or carbon evaporation ^14^.

Nowadays, commercial CLEM dishes or coverslips are routinely used for correlating fluorescence imaging of fixed or living cells with transmission EM (TEM) ^15,16^. Typical sample preparation for EM, i.e. by chemical fixation or high pressure freezing, includes a resin embedding step. Upon removal of the coverslip from the resin block, the region of interest (ROI) is located using the topology of the coordinate system that marks the block surface. For TEM imaging, the block is then trimmed so the sections containing a ROI can fit onto an EM grid. Regardless of the initial dimensions of the substrate, selecting the ROI usually entails the loss of surrounding areas, preventing the analysis of multiple cells if they were distributed across the full surface of the culture dish or coverslip.

In recent years, volume scanning electron microscopy (SEM) modalities have been used for CLEM on cultured cells. Besides offering access to large volumes, both serial block face SEM (SBEM) ^17^ and array tomography ^18,19^ also require block trimming before imaging and therefore suffer from the same limitations as TEM when utilized for CLEM. Focused ion beam SEM (FIB-SEM) ^20^ however can accommodate the imaging of large specimens without the need for trimming. In particular, multiple cultured cells grown on a Petri dish or coverslip can be imaged in a CLEM workflow, even when scattered across the full surface of the substrate ^21^. Despite this capability, CLEM has been performed one cell at a time and for a limited number of cells ^21,22,23,24^, because up to now, FIB-SEM microscopes lack automation procedures to acquire multiple sites without interruption.

In this paper, we introduce CLEM*Site*, a software prototype which automates serial FIB-SEM imaging of cells selected previously by fluorescence microscopy. We show that the automation is not only possible, but it also significantly reduces the number of required manual interventions during EM imaging. In addition to the automation process, we also describe the system of landmark correlations used to find targeted cells spread over the surface sample. Our software was tested in two types of CLEM experiments, each experiment type repeated twice. In the first type of experiment, for each session we selected around 25 cells from the same dish, each cell belonging to a different phenotype. In the second experiment, the same amount of cells was selected randomly, this time with only one phenotype present in the dish. We collected a significant number of EM images from multiple cells, which allowed us to conduct morphometric analysis on different phenotypes.

## Results

### Presentation

By following the logical workflow of a CLEM experiment, CLEM*Site* was designed modularly (Fig. 1a). The first module, CLEM*Site*-LM, is a stand-alone application to process sets of images acquired by light and fluorescence microscopy. CLEM*Site*-LM primarily extracts stage coordinates of target cells and their associated landmarks. The second module, CLEM*Site*-EM, is divided into 3 components that assist with automation: *Navigator* to find and precisely navigate to the targets, *Multisite* to trigger a FIB-SEM run on each position, resulting in a stack of serial images of the corresponding ROI, and *Run Checker* to supervise operations during each acquisition. To control the FIB-SEM microscope, CLEM*Site*-EM interfaces commercial software (*SmartSEM* and *ZEISS Atlas 5* from *Carl Zeiss Microscopy GmbH*).

**Figure 1.**
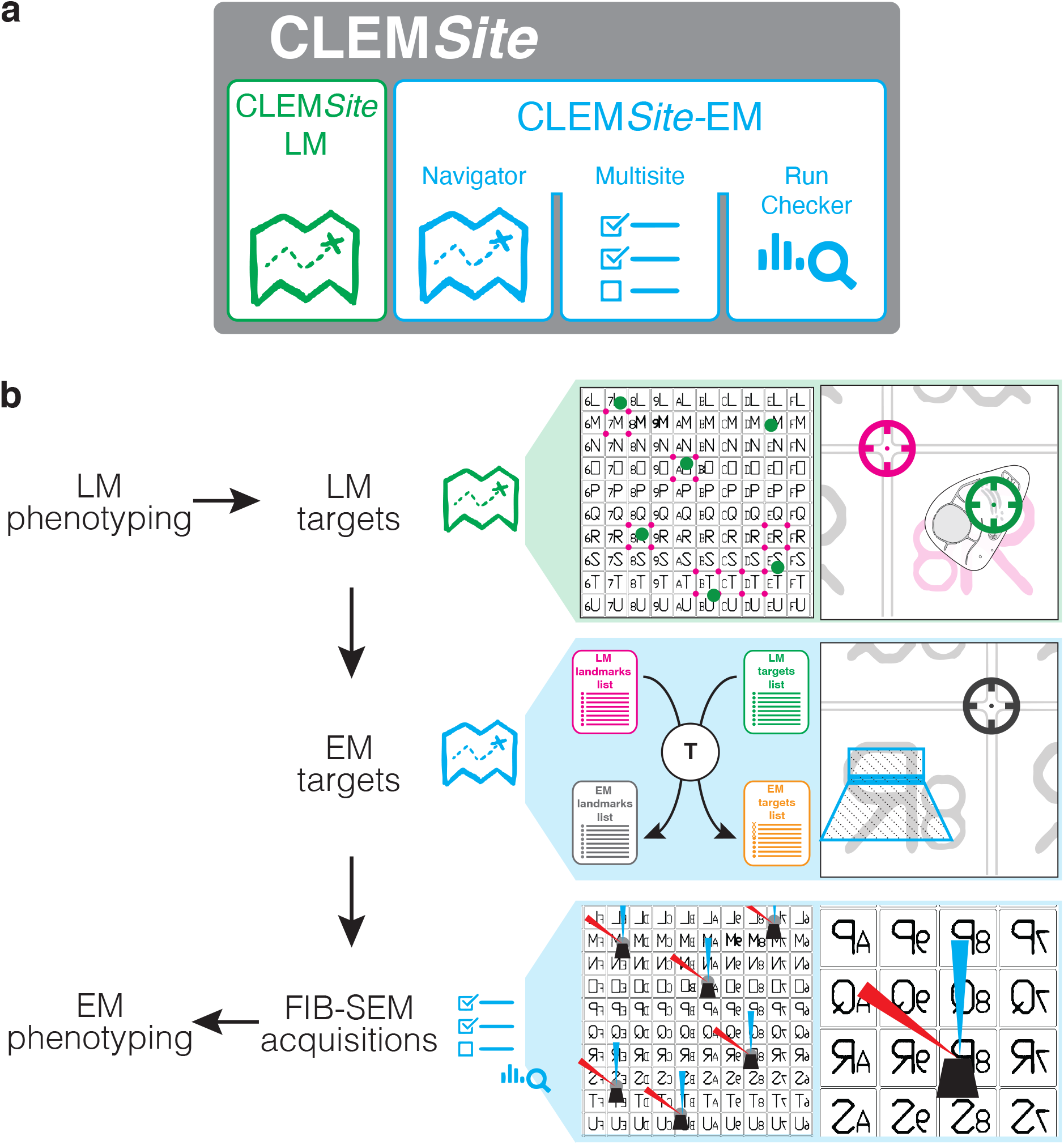
Schematic representation of the correlative light and electron microscopy software CLEM*Site*. (a) Overview of the different elements of CLEM*Site*, CLEM*Site*-LM and CLEM*Site*-EM. CLEM*Site*-EM is divided into 3 modules: the *Navigator*, which allows to store and move to different positions in the SEM, then *Multisite*, which drives the FIB-SEM acquisitions and the *Run Checker*, which controls and reports on the FIB-SEM runs. (b) Workflow for the automated acquisition of multiple correlated datasets. Light microscopy is performed, finding LM targets and recording their corresponding landmarks using CLEM*Site*-LM Inside the FIB-SEM, EM targets are predicted using the *Navigator* and acquired using *Multisite* and *Run Checker*. The acquired data is finally analysed to characterize different phenotypes.

### Correlation strategy

The correlation strategy applies transformations to translate cell positions (microscope stage coordinates) from LM into cell positions of the FIB-SEM (Fig.1b). At the light microscope, cells of interest can be selected either by manually screening or by using more assisted pipelines, such as the ones described in the application examples below. In our experiments, the Golgi apparatus morphology was used to select cells by means of an automated phenotypic screen. At each position where a cell of interest is identified for downstream CLEM analysis, a light microscopy acquisition job is programmed to collect a set of images. The first set comprises one fluorescence image at low magnification (using a 10x objective) (Fig.2a), and one reflected light image of the same field of view revealing the grid pattern (Fig.2b). The target area, which can be a cell or more precisely a subcellular region (e.g. the center of mass of the Golgi apparatus, Fig.2a) is placed at the image center. With a target centered, the stage coordinates are recorded for subsequent use in the correlation.

**Figure 2.**
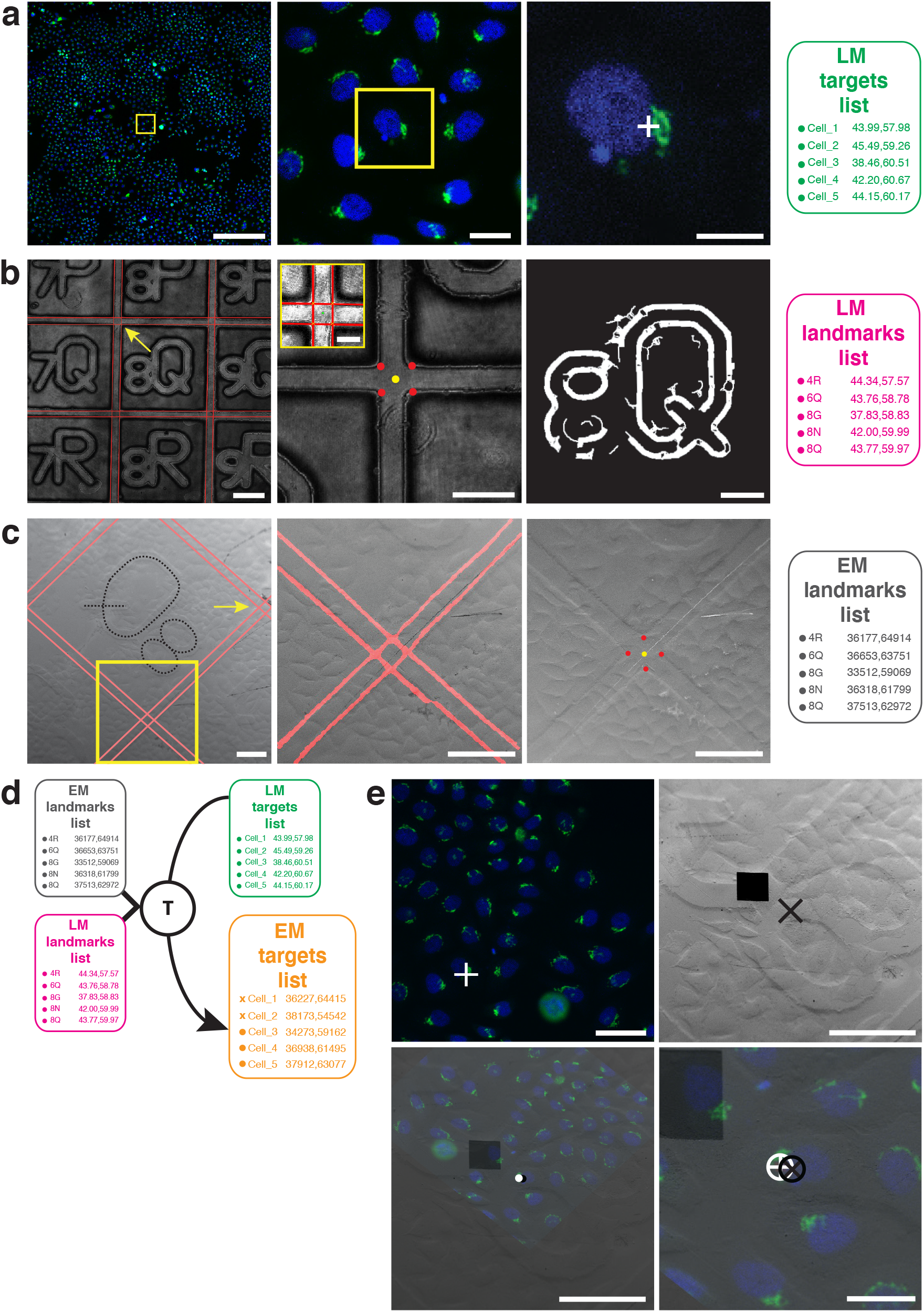
Coordinate system mapping and automatic detection for the correlation strategy. When a cell of interest is selected, for example to target the Golgi apparatus center of mass (a, white cross), the low magnification fluorescence (a, left image) and reflected light images (b, left image) are stored, and later loaded into CLEM*Site*-LM. This module extracts the stage coordinates in micrometers from each image metadata to build a list of targets (*LM targets list*) corresponding to the fluorescence , and another list of landmarks corresponding to the reflected light images (*LM landmarks list*) (b, middle image including inset). In the reflected light images, where the grid bars cross (red lines, band inset), the corresponding detected edges are converted to lines and mark 4 points (b, red dots), that are used to determine the center point (yellow dot). By convention the top left corner (yellow arrow) is named by associating its unique center point (yellow dot) with the alphanumeric identifier imprinted onto the glass dish bottom, which is automatically detected (b, right image). In the FIB-SEM, an image is taken by the *Navigator* module (c, left image), and the grid bar crossings are detected (c, middle image) to calculate the center point (c, right image and yellow dot), this process continues at each predicted landmark to give a list of landmarks (*EM landmarks list*). (d) A transformation is computed to register together the *LM list* and *EM landmarks list*, which is then applied to the *LM targets list* to predict the *EM targets list* across the sample at the FIB-SEM. (e) FM (top left) and SEM (top right) images were superimposed manually using the cell contours. For this the FM images were flipped, rotated and scaled (bottom left). The position of the LM target (white cross) is then compared with the predicted target in the SEM (black cross). A final targeting accuracy of 5± 3 *µm* (RMSD over n=10) was estimated by overlaying the SEM and LM images (bottom right). Scale bars: (a) 200 *µm*, 25 *µm*, 25 *µm*; (b) 200 *µm*, 100 *µm*, inset: 50 *µm*, 100 *µm*; (c) all 100 *µm*; (e) 50 *µm*, 100 *µm*, 100 *µm*, 25 *µm*.

All images are then loaded to CLEM*Site*-LM. The first step of the software is to automatically extract landmarks that will be used as references to register the stage coordinates coming from LM and EM images. The grid pattern imprinted on the bottom of the culture dish is a convenient coordinate system for registration. As the screened cells are typically distributed across the whole surface of the CLEM dish, a map of local landmarks is built from multiple sparse images of the grid.

Since the bars constituting the grid are relatively thick at 40 micrometers wide, the center of their intersections is used as a fiducial marker. In CLEM*Site*-LM, these centers are identified by a line detection algorithm, which is applied on the reflected light images to find the lines present at the grid bar edges. At the grid bar crossings, the detected grid bar edges intersect in 4 points, the centroid of which is used to mark the center of each grid bar crossings (Fig. 2b, and Supplementary Fig. 1). This center point is saved in stage coordinates as a landmark. Since each grid square is already imprinted with a unique combination of alphanumeric characters, each calculated center point is labelled using this existing identifier. Identification of the corresponding alphanumeric set of characters in reflected light images is performed by a VGG16 based convolutional neural network (CNN) ^25^ (Fig.2b). The CNN was trained with a combination of synthetic and manually annotated light microscopy images.

The last step in CLEM*Site*-LM, is to obtain a second collection containing the centroid stage coordinates of the target structures (e.g. the Golgi apparatus, Fig. 2a). In our experiments, since our target cells are centered on the image, stage coordinates are extracted directly from the image metadata.

After sample preparation for EM, removal of the coverslip and coating with a thin layer of gold, samples are transferred to the FIB-SEM chamber, where they are left to equilibrate for one day before starting the experiment. The next day, the examined sample is positioned for optimal visualization of the grid (see *Methods, Correlation in Electron Microscopy*). At the beginning, CLEM*Site*-EM requires an image from a random initial position of the sample surface to be used as a calibration step. The *Navigator* module prompts the user to indicate which grid square (identified by the alphanumeric identifier) is in the SEM image and in which orientation. The landmarks are then detected by the same line detector used by CLEM*Site*-LM. As a fail-safe, landmarks can also be manually identified by clicking over them.

Based on the manufacturer’s known grid layout and four landmarks, the software creates a linear model that represents a simple quadratic lattice to predict the position of all landmarks in stage coordinates. This preliminary model-based prediction of landmark positions has a targeting accuracy of approximately 5 ±20 *µm* (measured as RMSE, root mean square error) which is insufficient for precise localization of the cell and therefore requires additional refinements. This involves obtaining more landmarks along the surface sample. Thus, at each predicted landmark, an SEM image is automatically taken and a convolutional neural network (U-net ^26^) is used to compute the probability of each image pixel to belong to a grid bar edge (Fig. 2c). The line detector is applied again to the resulting grid bar edges to give the center point. This process is repeated throughout the sample surface to find and associate each individual landmark identified previously in light microscopy images.

When enough landmarks are collected, an affine 2D transformation is computed to register the landmarks from LM and EM. The transformation is applied to all LM stage coordinates of target cells to predict their position in SEM stage coordinates at the surface of the resin block (Fig.2d). When all 4 experiments are taken into consideration, this global transformation reduced the error in target accuracy down to 13 ± 6 *µm*. If the grid pattern is sharp and the block surface does not present any defects such as cracks, scratches or dust, grid edges are detected perfectly and the center point of the landmark can be calculated with higher accuracy (Supplementary Fig. 2). In our case, we had two of such experiments, reaaching a global targeting accuracy (RMSE) of 8± 5 *µm*.

To further increase the targeting accuracy, a local transformation delineates the third and final targeting refinement. It is calculated before each FIB-SEM acquisition, using only the landmarks in close proximity to the target (a total of 8 landmarks falling in a radius of 1200 µm). By applying this local refinement, we obtained a final targeting accuracy of 8± 4 *µm* for all the experiments (average of n = 10 cells per experiment over N = 4 experiments), or of 5 ± 3 *µm* with the pristine blocks (average of n = 10 cells per experiment over N = 2 experiments). These results were validated by registering manually the fluorescence image and the SEM view of the sample surface in the predicted position (Fig. 2e, Supplementary Table 1).

Thus, with our experiments, we exemplify how it is possible to perform an automated detection and registration of landmarks from both LM and SEM imaging modalities, which can lead to a final correlation with an accuracy of targeting close to 5 µm. Besides, the correlation can be performed over relatively large sampling areas: in the experiments a surface region of approximately 8 *×* 8 *mm*^2^ was completely mapped.

### Automation of FIB-SEM imaging of multiple cells

Once the correlations between cell positions in light and electron microscopy have been determined, the *Multisite* module of CLEM*Site*-EM executes the FIB steps of our automation workflow. These are automated localization of the coincidence point (Fig. 3a); milling of the trench to expose the imaging surface and the automated detection of this trench to ensure a well-positioned imaging field of view (FOV) (Fig. 3b); automated detection of image features in this imaging surface for the initial autofocus and autostigmation (*AFAS*) (Fig. 3c); and finally the stack acquisition (Fig. 3d). These four steps are executed sequentially for all targets (Fig. 3e).

**Figure 3.**
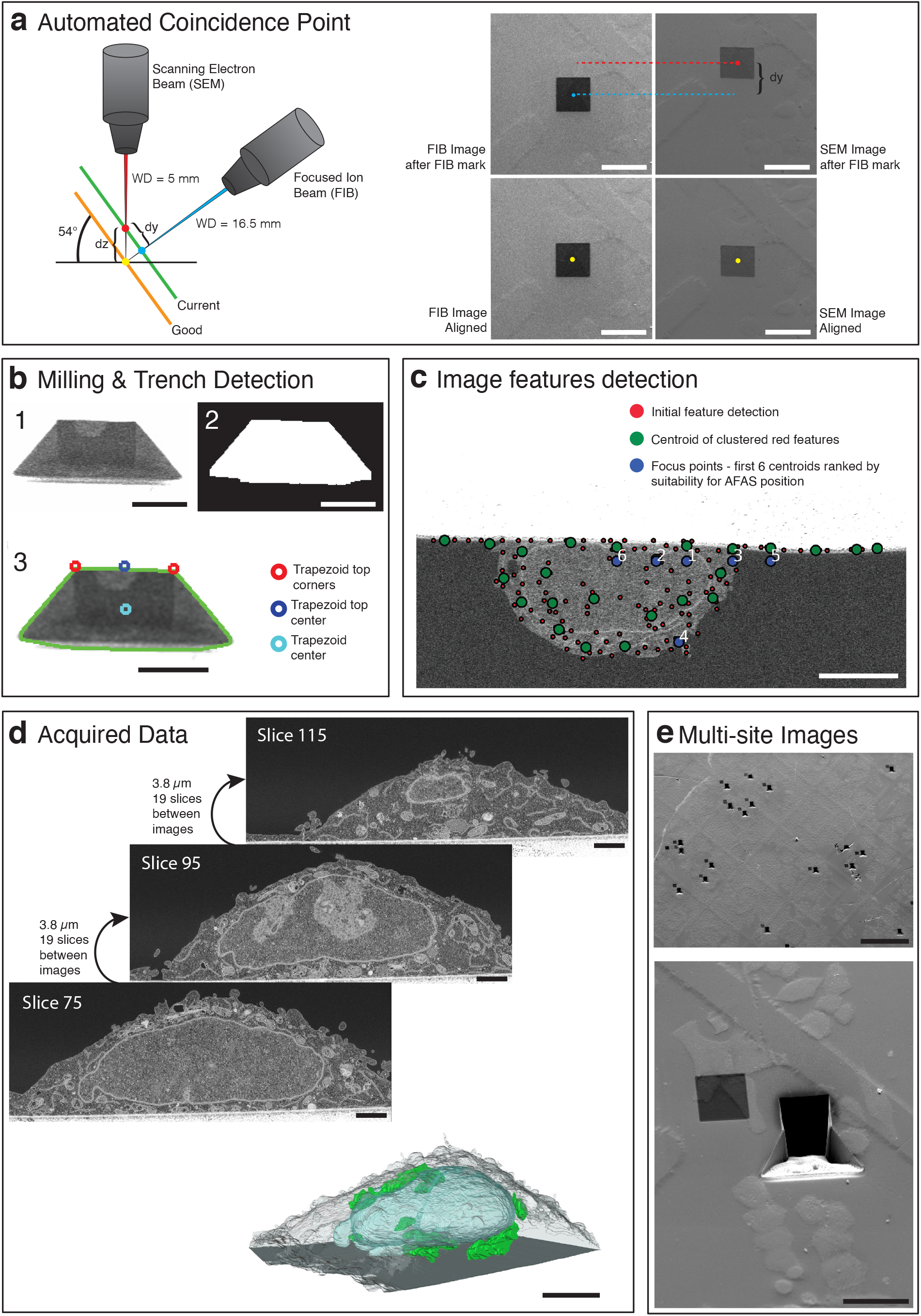
Automation of FIB-SEM workflow for multiple targets. (a) Automated Coincidence Point routine is illustrated schematically. When not tuned, the two beams are usually pointing at different positions of the sample surface (green plane, blue point for FIB center, red point for SEM center). The orange plane below, shows the case where the ideal position (yellow point) is achieved with respect to both FIB and SEM beams. In the software routine, a square is sputtered with the ion beam on the sample surface. The offset between the two beams is calculated based on the difference between the center of the sputtered mark in the SEM and FIB images (dy, distance between red and blue positions in the green plane). The z height (dz) of the stage is then corrected, and a further refinement using the SEM beam shift is performed by calculating the translation of the square mark between FIB (50 pA image) and SEM images. (b) Milling & Trench Detection: (1) After finding the coincidence point, a trench is milled to expose a cross-section at the region of interest. (2) The trench is detected to accurately position the field of view. First, a three level thresholding is applied to the image, followed by the detection of the biggest connected component that fits a trapezoid shape. From the final binary shape, boundaries of the trapezoid are found (3): the top corners (red circles), the trapezoid top center (blue circle) and the trapezoid center (light blue circle). (c) Image features detection: The image of the cross-section surface is analyzed and scored for the best focus positions to perform autofocus and autostigmatism. Features inside the image are found by using Harris corner detection and a variance map. The initial features (red points) highlight the high contrast and complex areas of the imaging surface which usually cluster on cellular structures. Features are clustered and their centroids (green dots) are then filtered and prioritized to detect the first 6 ones suitable for *AFAS* (blue points). (d) Acquired data: Images are acquired at 200 nm intervals (in z) throughout the Golgi apparatus region. The resulting stack is used for 3D render and quantifications. (e) Multi-site Images: Result of an experiment, where multiple targets had been acquired automatically across the full surface of the sample. Scale bars: (a) all 50 *µm*; (b) all 25 *µm*; (c) 5 *µm*; (d) slices all 2 *µm*, model 5 *µm*; (e) 500 *µm*, 50 *µm*.

The sample is positioned at the target coordinates of the first cell, and the *Multisite* module performs the coincidence point alignment of both the electron and ion beams (Fig. 3a and Supplementary Fig. 3a). To preserve the target, the sample is drifted 50 *µm* in x. The working distance is checked by autofocus and adjusted by the z-movement of the stage. A square fiducial area (20×20 *µm*^2^) is then created at the surface of the block by FIB sputtering at high current (7 nA). This square is then imaged by FIB and SEM sequentially (using the SE detector). The offset (in the y direction) between the center of the sputtered square (i.e. the focus point of the ion beam) and the center of the e-beam image is then utilized to calculate the z-offset by applying a trigonometric relation (Supplementary Fig. 3). A further refinement is achieved by cross-correlating images of the sputtered mark captured using the FIB (imaging current, 50 pA) and SEM modes. The measured difference in micrometers is then applied to the SEM beam shift to correct the FOV position.

Following the automated coincidence point alignment, the software proceeds with estimating the position of the target cell using the local transformation based on the closest landmarks as described above. After moving back to the estimated position, the software automatically triggers *ZEISS Atlas 5* to mill a trench, which exposes a cross-section orthogonal to the surface of the block. When the milling is finished, an SEM image is taken with the ESB detector and the trapezoid shape of the trench is detected using thresholding and shape recognition (Fig. 3b, Supplementary Fig. 3b).

The center of this shape is used as a reference to position the FOV to be imaged during volume acquisition. The FOV is magnified from a 305 *µm* by 305 *µm* to a 36.4 *µm* by 36.4 *µm* surface area and an image of the cross-section taken (Fig. 3c). A feature detector (Harris Corner detector ^29^) is applied to this image to identify salient points with high contrast and complex pixel neighborhoods. Such point features usually cluster around complex cellular structures, therefore they can be clustered using k-means. The k-means centroids are additionally filtered and prioritized by higher variance, high entropy and for their proximity to the center of the image. The first element in the filtered list can thus be stored for the subsequent application of autofocus and autostigmation procedures (*AFAS*) (Supplementary Fig. 3c). In the absence of a cell on the cross-section, the *AFAS* function is automatically targeted to the edge between the cross-section and the surface of the block. An image stack is then acquired (Fig. 3d) at a given regime (z resolution) as defined in the initial setup of the experiment. Note that whilst every cell of one run can be acquired with the same recipe (as defined in *ZEISS Atlas 5* : sample preparation, total volume to be acquired, slice thickness and FIB currents applied at each step), CLEM*Site*-EM also offers individual definition of recipes, allowing a per cell adaptation of the shape or volume (Supplementary Fig. 4a).

The *Run Checker* module of CLEM*Site*-EM (Supplementary Fig.4b) supervises each stack acquisition and corrects the position of the FOV if an image drift occurs in the y-direction. The drift is computed by using ASIFT ^30^ point feature correspondences, which are optimally filtered using RANSAC ^31^. When a drift is detected, the next image is corrected accordingly by adjusting the SEM beam shift. *Run Checker* also continuously monitors the run for the periodic autofocus and stigmatism. For each image acquired, a Vollath’s autocorrelation and a Laplacian metric ^32^ are used to measure respectively the quality of focus and stigmatism. When these values differ more than 25% between two consecutive slices, in addition to a warning UI message, an e-mail is automatically sent to the user, who can then decide to interfere and correct the drift manually.

After completion of one volume acquisition, CLEM*Site*-EM restores the original microscope conditions, drives the stage to the next target cell (using the *Navigator* module) and starts a new FIB-SEM run (*Multisite* module). This process is repeated until all targets are acquired (Fig. 3e). When the Gallium source is no longer producing a coherent ion beam, the FIB interrupts the current run. Upon reheating of the Gallium source, the run is then manually resumed to proceed with the next cells. For a typical FIB-SEM acquisition recipe at our microscope (as outlined below in case study 1), 15 to 20 consecutive cells can be acquired before it becomes necessary to reheat. CLEM*Site*, thus provides a unique solution for automated targeted 3D acquisition of multiple cells previously identified by light microscopy.

### Applications

We illustrate CLEM*Site*’s capabilities with two applications. In the first, the Golgi apparatus morphology of HeLa cells is perturbed with siRNA knockdowns by adapting a previously described solid phase reverse transfection protocol ^35^, where several siRNA knockdowns can be performed in a single experiment. This approach represents an efficient screening tool to identify specific genes involved in Golgi apparatus morphology.

In the second application, we illustrate a follow up of this screen, where morphological perturbations of the Golgi apparatus are further evaluated by focusing on one of the siRNA treatments, i.e. knocking down the COPB1 gene expression. This treatment was chosen based on its prominent phenotype. Variable transfection efficiency leads to a heterogeneous distribution of the phenotypes. We address this heterogeneity with our CLEM approach in which the target cells are selected according to their phenotype as visible by fluorescence microscopy. Using such a phenotype-enriched selection of cells enables us to collect sufficient data for a morphometric evaluation at the EM level.

#### Case study 1: Integrated multiple knockdown CLEM screen

Organelle morphologies can be observed by fluorescence light microscopy and used as a proxy to identify which genes are involved in various cellular functions. Previous experiments showed how the Golgi apparatus organization can be studied by tagging GalNAcT2, a resident enzyme of the Golgi apparatus, with a fluorescent protein ^33,34^. For efficiently screening the effects of different knockdowns, we have adapted an integrated experimental approach based on solid phase reverse transfection ^35^. By depositing drops of siRNA transfection mix, multiple treatments are distributed as an array at the surface of one single gridded culture dish. With such a layout, up to 32 spots can be deposited (Fig. 4a).

**Figure 4:**
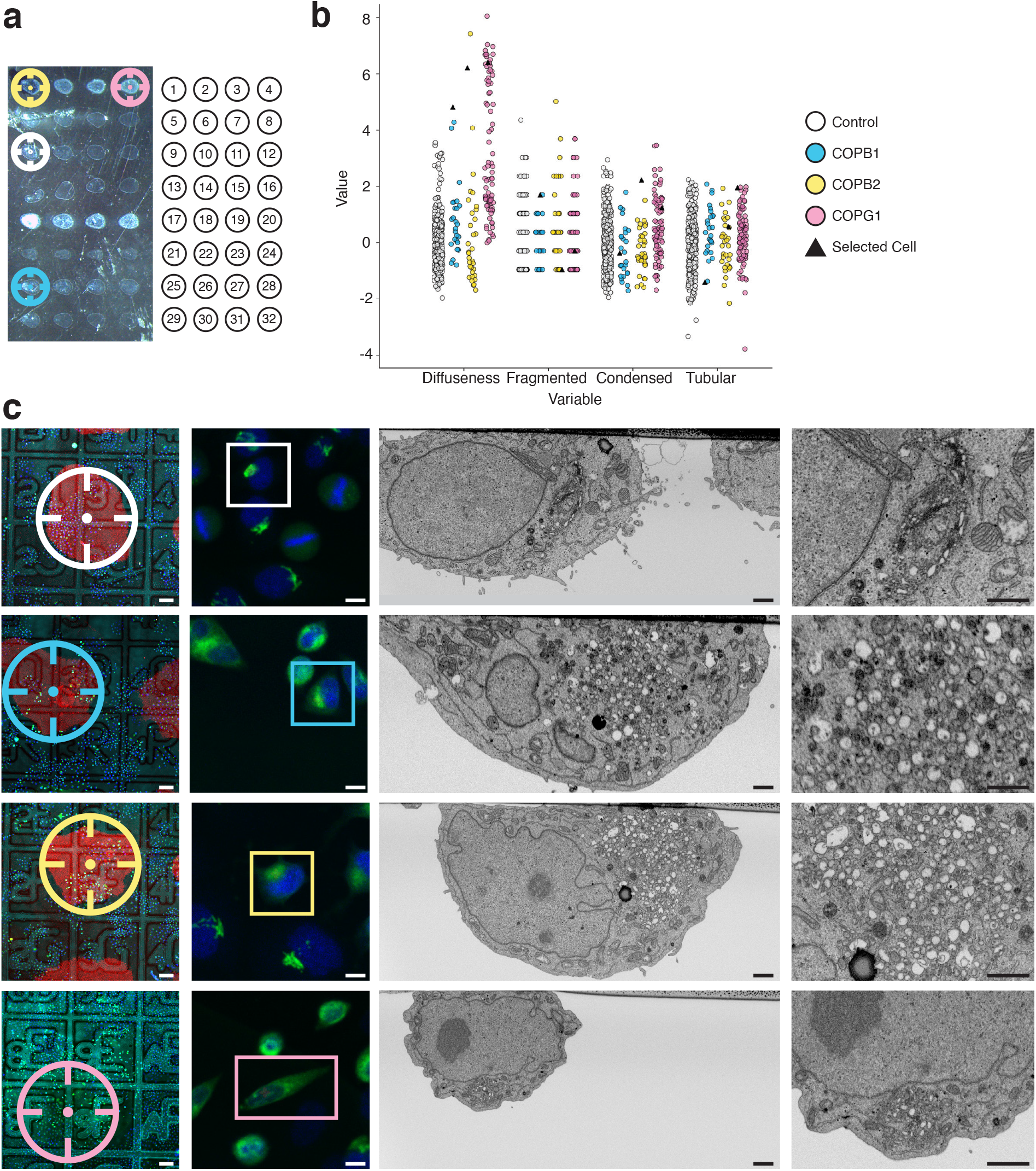
Automated screen of 14 siRNAs after 72h solid phase transfection knockdown. (a) Transmitted light image of one Petri dish with the 32 siRNA spots (left), where each siRNAs transfection mix is placed in the culture dish following a definite arrangement, see Supplementary Table 2 for further details (right). (b) Morphological features of the Golgi apparatus scoring tubularity, diffuseness, fragmentation and condensation for COPB1, COPB2, COPG1 in comparison to control. Values of each feature are normalized with respect to the mean of the control, n = 2985. During the light microscopy workflow, cells transfected with COP siRNAs display a phenotype that can be identified because of their high value in diffuseness. As an example, we selected one cell of each COP related siRNA (black triangles), to display in (c) the final result of the correlative experiment. (c) Selected correlated cells control, COPB1, COPB2 and COPG1 (top to bottom): overview merged fluorescent, reflected light image and image of the siRNA spot (left), fluorescent image of selected cell (second from the left), cross-section through selected cell in the region of the Golgi apparatus acquired automatically with the FIB-SEM (two images on the right). Scale bars: (c) left to right, 100 *µm*, 10 *µm*, 1 *µm*, 1 *µm*.

After a 72 hours incubation period, the cells on each siRNA spot are automatically imaged by confocal fluorescence microscopy. For this, four fields of view in each treatment spot are imaged with a 10x objective. The position of these fields is generated systematically using a matrix pattern. The resulting fluorescence images are then processed in CellProfiler ^36^, where the nuclei (DAPI channel) and a total of four features associated to the Golgi apparatus (GFP channel) from individual cells are extracted. Upon perturbation of the secretory pathway, the Golgi apparatus morphology can display a variety of phenotypes ^37^ which we classified into four typical appearance categories: fragmented, diffuse, tubular and condensed (Supplementary Fig. 5a). We designed the four features to score each one of such morphologies individually (fragmentation, diffuseness, tubularity and condensation) to measure the impact of each siRNA treatment (Supplementary Fig. 5b). Thus, a high score on one of the features serves as an indicator of the presence of the phenotype.

For this proof of concept experiment, the expression of 14 genes was challenged (Supplementary table 2). The most striking effects were observed when perturbing the expression of subunits of the COP1 complex, associated with non-clathrin coated vesicles (Fig. 4b,c). For the 3 subunits tested (COPB1, COPB2 and COPG1), a significant number of cells started to display a diffuse GalNAcT2-GFP signal, as visible by fluorescence microscopy after 72 h of treatment (Fig.4b,c). Under these experimental conditions, the other gene knock downs did not display noticeable phenotypes (Supplementary Fig. 5b), likely because the sample size was not big enough to detect subtle variations in the Golgi morphology.

Applying our automated CLEM workflow, we selected 2 to 3 cells per condition for further ultrastructural analysis by FIB-SEM. A total of 34 cells were automatically targeted (plus 2 control cells acquired manually) and acquired across 3 runs. For treatments with siRNA perturbing the expression of subunits of the COP1 complex, the cells were chosen from the pool that displayed the highest diffuseness score (Fig.4b, cells highlighted as triangles on the plot), a pool that was clearly distinguishable from the control condition. For other genes, even though the image analysis did not reveal any outstanding sub-population, we picked randomly between the cells displaying the highest scores associated to the expected phenotype, as hypothesized from previous experiments (Supplementary Fig. 5b, selected cells highlighted as triangles).

At the EM level, 5 out of the 6 cells treated with COPB siRNAs with a diffuse phenotype, displayed total disruption of the Golgi stack, which would normally display 3 to 4 closely associated cisternae. Instead, the region with enriched GalNAcT2-GFP fluorescence signal was filled with numerous vesicles (50 to 300 nm in diameter), suggesting comprehensive disassembly of the Golgi stacks upon knocking down the COPB1, COPB2 or COPG1 genes (as observed in the COPB1 of Fig.4c). For the remaining cell, a mixture of Golgi stacks and vesicles was observed.

The selected cells from the other siRNA treatments (Supplementary table 2) were also imaged by FIB-SEM in order to detect any subtle perturbations of the Golgi morphology at the ultrastructural level. For each condition tested though, Golgi apparatus was visible and stereological analysis ^38^ of stack composition or stack volume did not reveal any differences with respect to the control (Supplementary Fig. 5c).

Altogether, this experiment shows that our software can be utilized to screen for cellular and subcellular phenotypes in a large-scale CLEM experiment. When used in an integrated experiment with different siRNA treatments, CLEM*Site* enables automated and fast screening for protein knockdown effects on the fine ultrastructure of the Golgi apparatus.

#### Case Study 2: Screening for phenotypes

Specific gene knockdowns lead to perturbed phenotypes of the Golgi apparatus. As shown in the previous experiment, a striking phenotypic change occurs when cells are treated with siRNAs targeting subunits of the COP1 complex. Integrated screens with several treatments provide a reduced surface area where cells are exposed to siRNA. This in turn limits the number of phenotypic cells accessible for each condition. Therefore, we performed a secondary experiment, where the entire cell population of a culture dish was exposed to the treatment. We focused on a COPB1 siRNA treatment by liquid phase transfection and evaluated after 48 hours of incubation. Even though a larger number of cells displayed a diffuse phenotype under these conditions, the observed phenotypic diversity justified the use of CLEM to perform an ultrastructural analysis on the most perturbed cells.

As described above, a measure of cytoplasm fluorescence intensity levels was used as a score to select the diffuse phenotype. By defining a threshold on this score, all cells with a high value of cytoplasmic diffusion were selected and then the diffusion phenotype validated manually for each cell using a customized Jupyter notebook (see methods). Using adaptive feedback microscopy ^39^, the identified target cells were automatically re-imaged on the LM, acquiring the image sets necessary for the correlation (reflected light and confocal fluorescence at 10x magnification, see *“Correlation Strategy”*). Higher magnification z-stacks of the cell and Golgi apparatus were also acquired with the 40x objective (zoom factor x4) to document the spatial distribution of the organelles. The 3D information acquired here was valuable, for example in order to be registered to the 3D FIB-SEM volumes ^23^.

In the next step, the set of LM images was processed as described previously, to establish a list of LM landmarks and a precise list of target cell locations. The cells were prepared for electron microscopy and transferred to the FIB-SEM where CLEM*Site* autonomously acquired image stacks at each target location. In the example shown in Fig.5, the LM screen resulted in the selection of 90 cells. It is important to keep this initial number high in order to compensate for the loss of targets when progressing downstream in the workflow. The first selection removes the cells that are too close to each other (less than 150 *µm*) or that are on regions damaged during sample preparation (resin defects, scratches at the surface of the block) (see methods “Correlation in electron microscopy”). Following this filtering step, a final selection of 35 cells was acquired as FIB-SEM stacks, 25 of which were of sufficient quality for analyzing the fine morphology of the Golgi apparatus. Altogether, 25 cell volumes were acquired and analyzed (Fig. 5a) over an automated run that lasted 8 days, including one stop for manual reheating of the gallium source. Note that these cells were distributed across a 40 *mm*^2^ surface area with a maximum distance of 8.2 *mm* between cells. Our program fully automatically and efficiently performs correlations between fluorescence microscopy and FIB-SEM data. As an example, the rendered segmentation of the FIB-SEM volume perfectly recapitulates the cell morphology seen in FM (Fig. 5b), demonstrating the accuracy of the correlation. The resolution of the FIB-SEM images is sufficient to analyze ultrastructural details of numerous cells. In our case (COPB1 knockdown) we could reveal how the Golgi complex transitions from a stacked organization to an accumulation of vesicles (Fig.5c).

**Figure 5:**
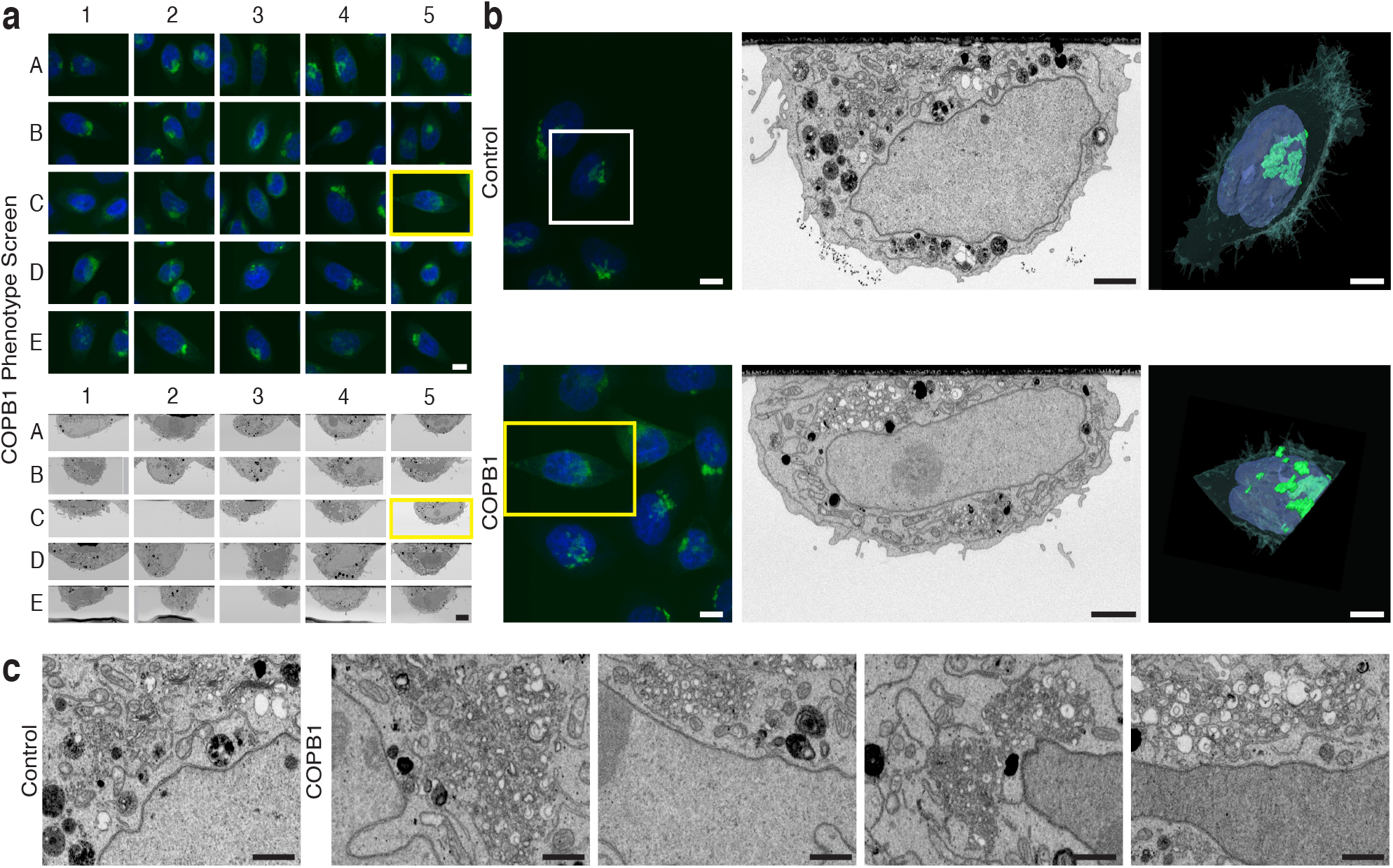
Automated screen on COPB1 cells in light and electron microscopy 48 hours after liquid phase transfection knockdown: (a) Overview of 25 selected cells in a screen for COPB1 knockdown. Light microscopy images (GFP GalNAc-T2 Golgi apparatus and DAPI for the nucleus, top) and the corresponding electron microscopy images (bottom). (b) Top: Selected control cell (treated with XWNeg9 siRNA) in light microscopy (left), electron microscopy (middle) and a reconstructed model from the FIB-SEM stack (right). Bottom: Selected COPB1 cell (treated with COPB1 siRNA) in light microscopy (left), electron microscopy (middle) and a reconstructed model (right). (c) Detailed electron microscopy images of the Golgi apparatus region in a control cell (left) and four different variations of a disturbed Golgi apparatus in different selected cells of the COPB1 knockdown. Scale bars: (a) LM - 10 *µm*, EM - 5 *µm*, (b) left to right - 10 *µm*, 2 *µm*, 5 *µm*, (c) 1 *µm*.

## Discussion

Performing CLEM on cultured cells, with cells selected for their phenotype during the LM step, is an efficient way to achieve ultrastructural analysis on subpopulations. Because this selection is performed prior to the EM sample preparation, these CLEM workflows are commonly performed one cell at a time. Because of an unprecedented efficacy to interrogate the cell ultrastructure in 3D, volume SEM imaging ^17^ is gaining popularity in the life sciences. Volume SEM has been used in CLEM experiments to capture phenotypic cells in culture ^20,21,23,40^. In most cases, volume SEM and CLEM were combined to capture the 3D ultrastructure at the highest spatial resolution possible (isotropic for FIB-SEM). Yet, the resulting low imaging throughput, in combination with individual cell picking, previously rendered volume SEM impractical for ultrastructural screens. Only in rare cases several cells were analyzed in a single experiment^21^. Consequently, volume CLEM is rarely used for screening large populations of cells.

Using the novel workflow relying on the software we developed, it was shown that correlative imaging using FIB-SEM can acquire multiple targets within a single experiment (up to 30 over one week of acquisition) with full automation. Detection of local landmarks imprinted in the culture substrate enables automated correlation and targeting with higher accuracy than previously achieved. The detection algorithm we developed could be extrapolated to other customized dishes or commercial substrates for cell culture in SEM samples^41^. Another advantage of local landmarks for the correlation is that they mitigate the impact of sample surface defects or optical aberration across long distances. Thanks to the utilization of a FIB-SEM, nearly the whole sample surface is accessible, enabling the correlation of multiple cells. Especially in the case of highly distributed and distant rare events ^42^ the respective targets are still within reach. We demonstrate the workflow on commercial dishes with a usable surface on the order of 40 *mm*^2^, but much larger surface areas are possible. In fact, the limitation is dictated mainly by the dimensions and travel range of the microscope stage. With such potential, the other main feature of our software is the ability to trigger an autonomous acquisition in multiple sites in one microscopy session. This fully automated triggering has not previously been achieved on biological samples. This is made possible through the automation of key steps of the imaging pipeline, i.e. 1) setting the coincidence point of both ion and electron beams, 2) automated evaluation of the image quality and, 3) constant tracking of the sample position within the field of view of the microscope. In summary, all interactions with the microscope that are usually supervised by a human operator during acquisition, be it several hours or days, are automated.

One essential paradigm shift for increasing the acquisition throughput is the decision to decrease the resolution in the z-dimension, thus prioritizing the speed of acquisition and ultimately the total number of cells acquired in one run. For many of the morphological features used, low z resolution has been proven efficient to score phenotypic variability at the subcellular level ^2^. Here, images in one volume are acquired every 200 nm, a step size much larger than typically used for isotropic voxel acquisition (e.g. 4-8 nm resolutions). The resulting gain in speed is significant, leading to only 6 hours necessary to acquire one full cell (including the creation of the trench). This is in stark contrast to isotropic acquisitions that can take from days to weeks per adherent cultured cells ^43,44^. Extrapolating acquisition time to a screen of about 30 cells, our workflow can deliver results in 10 days compared to isovoxel acquisition regimes that would require more than 6 months of machine time. Therefore, CLEM*Site* is intended to be a screening tool for performing quantitative assessments of morphological variations. It is efficient to reveal rare phenotypes at the ultrastructural level, and increases the number of observations. Other acquisition regimes of FIB-SEM can be considered if higher resolutions are required, at the cost of a (much) lower throughput ^24,44^.

Capitalizing on the software’s ability to screen across the full surface of the dish, we demonstrate that multiple siRNA treatments can be performed in a single integrated CLEM experiment (by spotting siRNA onto the culture substrate). Provided that other treatment reagents can be bound to the culture substrate we anticipate that the same approach can be expanded to screening the effects of various drugs on subcellular morphologies. While we focus on enhancing existing hardware with targeting and automation abilities in this work, the next challenge is to efficiently analyze the resulting large amount of data generated. So far, we are using the powerful tools brought by stereology. We think that following the same principles, especially when designing the sampling strategies ^38,45^, the manual assessment of subcellular morphologies will be replaced by applying state of the art computer vision, such as AI based semantic segmentation followed by morphometric analysis. Once these tools are readily available, CLEM*Site* will be endowed with even more power to support molecular cell biologists in morpho-functional studies.

## Acknowledgments

This work was supported by EMBL funds and by SFB1129. We thank the staff of the Electron Microscopy Core Facility and the staff of the Advanced Light Microscopy Facility at EMBL, Heidelberg. We thank Alexandre Laquerre and Ken Lagarec (*Fibics Incorporated*, Canada) for all the assistance in *ZEISS Atlas 5* software and *Carl Zeiss Microscopy GmbH* for support and feedback. We thank the Electron Microscopy Core Unit at the Max-Planck-Institute of Experimental Medicine Göttingen for access to their Crossbeam.

## Methods

### Cell culture

HeLa cells stably expressing GalNAc-T2-GFP 31 were maintained in Dulbecco’s Modified Eagle’s Medium (DMEM Dulbeccos Modifed Eagles Medium, Sigma Aldrich) culture medium containing 10% fetal calf serum (Gibco Life Technologies), 100 Units/ml penicillin (Gibco Life Technologies) and 100 *µg/ml* streptomycin (Gibco Life Technologies) and 2 mM L-Glutamine (Sigma Aldrich), incubated at 37 °C and 5% *CO*_2_. Cell selection was applied using 500 *µg/ml* Geneticin (G-418 sulfate, Gibco Life Technologies) for every passage of the cells. Cells were incubated on gridded MatTek dishes (P35G-2-14-C-GRID, MatTek corporation) with siRNA spots and incubated for 72 h in DMEM medium without phenol red.

### siRNAs

siRNAs targeting Golgi apparatus ^37^ morphology in this study were obtained from Ambion/ThermoFisher as Silencer Select reagents, please see Supplementary Table 2, for siRNA IDs and sequences.

### siRNA pre-screen and solid-phase reverse transfection

From an initial genome-wide screen for proteins affecting the secretory pathway ^37^, 143 siRNAs had an effect on the morphology of the Golgi apparatus. From them, 79 of the strongest phenotypes were selected. These were used in a pre-screen in order to find the most promising candidates for further CLEM experiments. 96-well plates (glass-bottom) were coated with siRNA transfection mixtures ^35^. Afterwards, HeLa Kyoto cells stably expressing GalNAc-T2-GFP (3400 cells/well) were seeded using an automated cell seeding device (Multidrop / ThermoFisher). Cells were imaged on a ScanR microscope (Olympus, UPlanSApo 20x 0.7 Ph2, DAPI, GFP and transmitted light). The plates contained control siRNAs for which the phenotype is well characterized on a light microscopy level: siRNA targeting COPB1, AURKB and KIF11 and, non-silencing negative control siRNA (XWNEG9). The 14 siRNAs showing the most prominent phenotypes were chosen for further CLEM experiments. Candidate selection was based on morphology of the Golgi apparatus, as visible from the fluorescent signal given by the GalNAc-T2-GFP. From collected images, morphological features were computed as explained below (Light Microscopy prescan and CellProfiler feature extraction). Selected siRNAs were spotted onto a gridded MatTek dish (P35G-2-14-C-GRID) using a contact spotter (ChipWriter Pro-Bio-Rad Laboratories) resulting in a layout of 4 × 8 spots 40. The mixture either contained oregon-green 488 gelatine (Thermo Fisher Scientific) or Alexa-494 gelatine (labeled with molecular probes protein labeling kit, Thermo Fisher Scientific) to make the spot boundaries visible. The array contained a total of 6 controls, as follows: 3 spots of negative control siRNA (XWNEG9), 2 spots of siRNA against AURKB and KIF11 (transfection control) and 1 spot of siRNA against COPB1. The other spots contained siRNA that target genes showing a Golgi phenotype after RNAi knockdown. 70000 cells per ml were seeded onto the spotted MatTek dishes and fixed with a light fixation (0.5% glutaraldehyde, 4% formaldehyde in 0.1 M PHEM) after 72 h of siRNA treatment. The observed transfection efficiency for a successful experiment is not uniform and oscillates between 40 to 70% within the spots.

### Liquid phase transfection for COPB1 knockdown

Liquid transfection with the siRNA (S3371) that is associated with the gene COPB1 was used in MatTek dishes (P35G-2-14-C-GRID) where cells were seeded at 70.000 cells/ml per dish. A standard protocol for transfection was used combining 3.3 µl of 30 *µM* siRNA with 1.5 µl of Lipofectamine 2000 (Invitrogen). Cells were examined by light microscopy 48 hours post-transfection.

### Fixation before light microscopy

Cells were fixed with a mixture of 4% formaldehyde and 0.5% glutaraldehyde (EM grade EMS) in 0.1 M PHEM Buffer (pH 6.9: 240 mM PIPES (Sigma), 100 mM Hepes (Biomol), 8 mM MgCl2 (Merck), 40 mM EGTA (Sigma)). A Ted Pella BioWave microwave with a temperature control unit (Pelco Biowave microwave with ColdSpot (Ted Pella Inc.)) was used to accelerate the fixation process to 14 min at 250 W. DAPI (1 µg/ml in 0,1M PHEM,Thermo Scientific) was applied to cells to stain the nucleus for a total of 10 minutes. To quench glutaraldehyde auto-fluorescence, cells were rinsed with 150 mM glycine (Merck) in PHEM buffer. Cells were left in the PHEM buffer for imaging.

### Light Microscopy

#### Light microscopy prescan and feature extraction

After the MatTek dish was mounted on the light microscopy (LM) stage (Leica SP5 MSA) the four corner spots of the siRNA array (Figure 4a) were located based on their green/red fluorescence using a 10x lens. At each corner, we used a python script to save the stage position from the microscope. After storing the positions of the 4 corners, the script generated a list of stage positions (2×2 sub-positions within each siRNA spot) that were loaded as positions onto the Leica Matrix Screener software. For the prescan images the following specifications were used: 10x objective, 680×680 pixels, zoom 6, FOV 258 *µm* x 258 *µm*, 4x averaging, sequential scan for excitations 405 nm (DAPI-labeled nuclei), 488 nm (GalNAc-T2-GFP), 594 nm (A594-labeled gelatine). Prior to each acquisition a software autofocus was performed on the DAPI signal. A CellProfiler image analysis pipeline (http://cellprofiler.org/releases/, version 2.2) was configured to segment nuclei based on the DAPI signal and then delimit cytoplasmic cell ROIs by radial dilation of each nuclear ROI. Within each cell ROI the GalNAc-T2-GFP signal was used to compute four intensity-independent features characterizing different typical alterations of Golgi morphology:

#### Diffuseness

Diffuseness was designed to characterize the fraction of GalNAc-T2-GFP signal dispersed in the cytoplasm. Given the image with the GalNAc-T2-GFP signal, with the cytoplasm already segmented for each cell the diffuseness of a cell is computed as the sum of all pixel values of the cell cytoplasm after a morphological grayscale opening, divided by the sum of all the pixel values of the cell cytoplasm. This value is high when the signal intensity of the Golgi is homogeneously distributed over the cell body.

#### Fragmentation

This feature was designed to characterize the number of seemingly unconnected Golgi structures. In some phenotypes, the Golgi apparatus is split into many pieces of variable size, with the biggest pieces being much smaller than a typical Golgi shape that would be observed in the negative control. Fragmentation was calculated by counting the number of separate connected components following a top hat morphological filter and Otsu thresholding ^46^.

#### Tubularity

Some phenotypes show high tubularity, also described as “enlarged” ^37^, where the Golgi apparatus had elongated cisternae running along the cytoplasm. Morphological grayscale openings of the GalNAc-T2-GFP signal are computed using structural elements in the form of a line at angle different orientations (0-180°). The difference between orientations that yielded maximum and minimum results of the filter are saved for each pixel. After this, the total value of elongation is computed as the sum of all values divided by the sum of the GFP intensity value for each cell.

#### Condensation

In other siRNA treatments, the Golgi was condensed in a smaller area, looking almost circular at the fluorescent microscope. Condensation was measured by the form factor, which is calculated using the formula 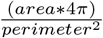 , after performing a top hat filter and an Otsu thresholding of the GFP intensity for each cell cytoplasm if the length of the outline from the thresholded signal delineates a perfect circle, it will match with the area of the enclosing circumference, providing the maximum value of 1.

#### Cell selection using a Jupyter notebook

Images from the previous step were stored in separate folders (one for each position where images were taken), and the CellProfiler pipeline applied to them. The output values for the features detected were stored in a comma separated value file. This file can be read by any statistical software to remove possible artefacts or undesired effects like dividing cells or cells too dim to be properly classified. This step, known as quality control (QC), is also useful to explore the results by analyzing the values of the controls with respect to treatments and doing exploratory analysis of our features. The QC and subsequent cell selection based on features was implemented in python and executed in a Jupyter Notebook ^47^. The main packages used together with the Jupyter environment are Numpy, Pandas ^48^ and Bokeh ^49^. Pandas was used to read the files from the CellProfiler pipeline and organize in tables the information associated with each cell (features and its original associated image). Bokeh and Holoviews enable us to provide interactive plots inside the Jupyter Notebooks that increase usability and speed up the process of manual cell selection. Once the features calculated in CellProfiler were loaded, cells expressing too little GalNAc-T2-GFP were rejected based on their integrated signal. Next, mitotic cells were rejected based on the coefficient of variation (CoV) of the DAPI signal, using the observation that, due to chromosome condensation, mitotic cells had a higher CoV than interphase cells within the segmented DAPI region. Finally, the PowerLogLogSlope feature ^50^, was used to remove potentially out of focus cells. After this, images of positive controls (AURKB, KIF11 and COPB1) were shown for qualitative assessment, and the experiment was continued if the three positive controls showed visible effects and the negative control was under standard conditions.

After the QC, which filtered out around 20 to 30% of the cells, the selection of cells for CLEM was displayed in a series of interactive plots. The plots include controls that allow cells to be selected individually. If the phenotype can be differentiated from controls by one of the main features (e.g. COPB1 by diffuseness, ACTR3 by condensation (Supplementary Fig. 5a)), cells can be selected inside a jitter plot based on their feature values. Some phenotypes could manifest synergy in the feature space, for example those that showed both fragmentation and tubularity. For this reason, t-SNE ^51^ can also be used to cluster populations naturally. In the t-SNE plot one can interactively select small clusters (as displayed in the Jupyter notebook).

After the coarse selection, small cropped preview images of each cell in the subpopulations for each gene are displayed. The user is prompted to individually confirm that the automatically selected cells exhibit the expected phenotype. The interface supports this through a yes/no button below each cell picture. A minimum of 42 cells were picked (3 per treatment in a total of 14 treatments, the transfection controls AURKB and KIF11 were excluded) for CLEM (Fig. 4, Supplementary Fig. 5a). The stage coordinates of the selected cells were saved and used to automatically guide the high-resolution imaging on the confocal microscope. For the liquid phase transfection experiment (Fig. 5), where only one phenotype was studied (COPB1), a total of 25 cells were selected, using only the property of diffuseness (Fig.5c) and selecting values higher than the control average.

#### High-resolution imaging light microscopy

For each of the cells selected by image analysis the following automated scan job pattern was triggered: (a) cell coordinates were passed to the microscope and the stage was positioned such that the selected cell was centered on the optical axis, (b) software autofocus on DAPI signal of the target cell using 40x objective, (c) high-resolution z-stack acquisition (9 slices, 10.1 *µm* range) with 40x objective, 512×512 pixels, zoom 5, FOV 77.5 *µm* x 77.5 *µm*, channels 405 nm/488 nm, (d) imaging of the spatial context of the cell including the etched coordinate system with the 10x objective, 1024⨯1024 pixel, zoom 1.2, FOV 1.29 *mm* ⨯ 1.29 *mm*, channels 405 nm/488 nm/594 nm fluorescence, transmitted/reflected light. Communication with the microscope software was implemented in python using a library of functions that communicate with the Leica Matrix Screener software ^39^. Two functions were used, one to move to the specific coordinate calculated in the previous step and another executed the acquisition of the described sequence of images, previously programmed using Leica LAS AF software (version 1.0.4, 2013).

### Correlation in Light microscopy

CLEM*Site* is a set of software tools developed in python and C# to support automated correlative light and electron microscopy (CLEM) (Fig. 1). The first of these tools, the CLEM*Site*-LM, was used to process the light microscopy images and extract landmarks from them (Fig. 2). The user provides a folder containing at least one image with two channels, one fluorescent with the GFP tagged organelle of interest and one showing the patterned glass bottom grid. Both images were acquired simultaneously at low magnification with a FOV that included a full square and patterned letter inside, usually 600 *µm*^2^. The grid pattern was acquired using reflected light (RL), modality of the confocal microscope. Images had a pixel size of 1.7 *µm*/pixel with a dimension of 1024×1024 pixels.

A script was created to rename the folder images to a more readable format and then, the set of folders was given to the application CLEM*Site*-LM. This application reads the RL image and applies the algorithm LOD (Line Orientation Detector) (Supplementary Fig. 1). LOD applied a series of preprocessing steps in which gradients are selected positively if they follow a line. Pixel orientations were weighted with neighboring pixels by convolution with line morphological operators for each possible angle orientation. A projection of the image onto one axis from 0 to 180 degrees, at a resolution of 1 degree, generated a 2D map where the main trend of a line inside the image could be detected by finding maxima using non maxima suppression. Once the most prominent lines were found, iterative refinement steps were applied to logically discard the lines which are not likely to belong to the grid, e.g. sets of lines not crossing orthogonally or not keeping approximately the expected measures given by the manufacturer of the glass bottom dishes.

Points resulting from calculating line intersections on the grid were associated with their corresponding alphanumeric identifier and used as landmarks. The LOD parameters are dependent only on resolution, where the number of neighbors was set to (k=12) and stroke size (stroke = 15) for 1024 × 1024 images. The parameters used for LM were between 0.06 to 0.075 for the Canny threshold, and a Laplacian filter was applied before in the presence of regular interference patterns or local contrast enhancement (CLAHE) when the brightness and contrast was unbalanced. The other provided parameters are the dimensions of the grid. In our case, we used MatTek grids, where the lattice is formed by a sequence of two lines spaced 40 *µm* followed by another space of 580 *µm*. We used as a landmark the center position of the small square formed by the intersection between two sets of perpendicular lines spaced 40 *µm*.

Each LM position in a map also requires a unique identifier. For landmarks, we conveniently named them after the two-character combination inside the nearest grid square (e.g. 4N). By convention, given a grid square with the inner pattern straightly oriented, the top left corner landmark is assigned (Fig. 2b). To automatically identify the characters on each grid square we trained a U-net using a mixture of real (20%, 1115 images) and synthetic (80%) binary images (128×128) of the identifier patterns combined with augmentation. The CNN architecture, implemented with Tensorflow, used 6 convolutions layers in a sequential manner followed by two dense layers. The loss function used was categorical cross-entropy that converged after 70 epochs. The prediction of the network was additionally validated using previously detected neighbors and the expected position of the landmark (e.g. 1A can be a neighbor of 1B, but not of 8B).

Detected landmarks are saved by the application in image coordinates. Since the stage coordinates of the optical microscope (saved in the metadata of the image) refer to the center of the image, the translation from pixel coordinates to stage coordinate can be obtained by simple addition after converting pixels to distances using the known image pixel size. Similarly, for each targeted cell, the difference in micrometers from the center of the image to the centroid of the object of interest was provided and converted to its respective stage coordinates.

### Electron microscopy

#### Electron microscopy sample processing

The entire EM processing was done using a Ted Pella Biowave Pro microwave. After samples were lightly fixed and imaged in the confocal microscope, they were heavily fixed with 2.5% glutaraldehyde (EMS), 4% formaldehyde (EMS) and 0.05% malachite green (Sigma) in 0.1 M PHEM (pH 6.9: 240 mM PIPES (Sigma), 100 mM Hepes (Biomol), 8 mM MgCl2 (Merck), 40 mM EGTA (Sigma)). The samples were then postfixed with 0.8% *K*_3_*Fe*(*CN*)_6_ (Merck), 1% *OsO*_4_ (Serva) in 0.1 M PHEM. The samples were stained successively with 1% tannic acid (EMS) and 1% uranyl acetate (Serva) to enhance the contrast. Samples were dehydrated in a graded ethanol series (30%, 50%, 75%, 90%, 2x 100%) and infiltrated in a graded series of Durcupan (25%, 50%, 75%, 90%, 2x 100%, Sigma) and polymerized in the oven at 60°C for 96 h.

#### Correlation in Electron Microscopy

The central disk of the MatTek dish was broken out using heat shock. The resin disk, containing the cells along with the imprint of the coordinate system on the surface, was mounted on SEM stubs (Agar Scientific) with a conductive carbon sticker (Plano). To reduce the amount of charging the samples were surrounded by silver paint (Ted Pella Inc.) and coated with gold for 180 seconds at 30 mA in a sputter coater (Quorum, Q150RS). The samples were introduced into the Crossbeam 540 (*Zeiss*) and positioned at 54 °. CLEM*Site* is interfacing *ZEISS Atlas 5* version 5.2.0.150 from *Fibics Incorporated* to navigate to the correct positions and to prepare the ROI for imaging. When scanning the surface of the sample with a scanning electron microscope (SEM), after detaching the glass from the resin, the imprinted grid pattern from the glass bottom dish and letter combination could be clearly observed.

To optimize the visualization of the gridded pattern, samples are rotated using the FIB-SEM stage to orient the grid at a 45°angle with respect to the SEM image (Fig. 2c). This ensures that both perpendicular orientations of the grid are efficiently detected when recording the secondary electrons which are best suited to visualize topological information. Cell contours could be visualized at higher accelerating voltages (5-10 kV) allowing signals to be detected from deeper regions inside the sample. However, cell visibility was sample dependent and individual cells could only be differentiated one from another at lower cell confluency.

Once all LM images were processed and the landmarks and targets stored in their respective files, we proceeded with the FIB-SEM acquisition. Our software connected to the microscope control in a client-server architecture, where the client streamed information and commands which were parsed, validated and then executed by a server. The server software relied on a dynamic library in .NET provided by *Fibics Incorporated* to control the microscope via *ZEISS Atlas 5*. When the microscope was ready, a first image of the surface was acquired (1.5kV, 700 pA) using the secondary electron detector (SESI) and sent to our client application. The reliability of the computational process was increased by having the user move to a grid square and indicate the combination of visible characters in that square. After the first image was computed, landmark references were calculated and mapped to an ideal coordinate system layout based on manufacturer measures (MatTek dishes: 560 *µm* x 560 *µm*, line width of 40 *µm* for SEM), and initializing a linear system to predict further positions within the grid. Afterwards the stage was moved to the approximate position of each grid square close to the regions of interest to be acquired.

When applied to SEM images, the LOD algorithm (used previously in LM) had a high failure rate (from 0.05% to 5%). In SEM images, grid lines are very often blurry or erased. Neural networks have proved to be very resilient to noise in classification and object detection ^52,53,54^. Based on those successes, we trained a U-Net network ^26^ to provide the probability mask where edges of a crossing could be found (Fig. 2c). Enough training data to optimize the network was provided with data from the LOD and used with the errors curated manually. Manual segmentation was performed using the corner shadows in around 100 difficult cases, when LOD failed. We extracted a total of 600 images and augmented them to 3000 images by variations in scaling, rotation, translation and intensity values. As preprocessing steps after augmentation, CLAHE (32×32 filter size), gaussian blur (sigma 1, 5×5 filter size) and normalization were applied. Processed images were then used to train a convolutional autoencoder based on U-Net, using binary cross-entropy as loss function, together with an Adam optimizer at a fixed learning rate of 1e-4. The network computed a probability map of the regions in the image that contained an edge belonging to a grid line. The last part of the previous LOD algorithm was adapted to find the peaks based on the maximum probability of lines and provided results in the form of image coordinates. Using the detection system based on CNNs the rate of failure in detection was reduced, with an average RMSE of detection originally of 12.66± 18.8 um to one of 6.23± 6 um respect ground truth (with n=149), a considerable improvement. The implementation of all the networks was carried out with Keras 2.0.8 and Tensorflow 1.3.

The described detection step was done for 30% of the grid corners, with a minimum of 20 grid corners required. The scanning of the surface sample (duration between 30-40 minutes) provided enough positions to generate a global registration with a targeting-accuracy of 5 to 20 *µm* throughout a surface area of 1 *cm*^2^. Since not all positions acquired in LM could be present in the current resin block (in several occasions the resin block broke into two different pieces), another CNN was trained to detect positions very close to the border or outside the sample, with a binary output (image valid or not). After the scan was finished, positions of targeted cells were predicted using a global affine transformation, which uses all available matched points between LM and SEM. All the information from mapping was stored in memory using a pandas dataframe for further usage and queries. If cells were closer than 150 *µm* one of them was removed, because the trench and acquisition of the first cell could interfere with the acquisition of the adjacent second cell.

The generated map between LM and SEM was used prior to each acquisition of a targeted position. First, it was used to move to the target cell and calculate the coincidence point there. After this, any landmark in a radius of 1200 *µm* close to the region of interest was imaged (usually resulting in four to eight neighboring landmarks) and stage coordinates were obtained again. Subsequently, for each predicted position an accuracy of 2 to 5 *µm* was achieved. This was precise enough to hit the organelle of interest.

In the current workflow, analyzing the images from light microscopy to extract the positions of the cells and the glass bottom grid took approximately 2 hours. This can be done in any computer at any time between the LM and EM session. In this step extra verification steps were added that help the researcher to validate the current cells selected in the light microscopy images. The corresponding map of the grid in the FIB-SEM in the resin block is acquired in approximately 3 hours. The initial setup for the first volume acquisition takes around 30 minutes, with minimal user input (brightness and contrast of detectors). After the first cell acquisition starts successfully, the microscope can run autonomously until the FIB-Gallium source has reached its limits and needs to be reheated.

#### Automation of the Focused Ion Beam - Scanning Electron Microscopy acquisition

To align the electron-beam and the ion-beam, we used an automatic coincidence point procedure to match both (Supplementary Fig. 3a). First a square was burned onto the surface of the sample using the FIB. The geometrical relation between the two beams was used to move the stage to the coincidence point resulting into both beams hitting the same spot on the sample surface (Fig.3a). After this, a trench was milled in front of the ROI at a FIB current of 15 nA and a 20 *µm* depth. Polishing was performed at 3 nA. In this way a cross-section through the ROI was exposed and the polygonal shape of the trench was detected (Fig. 3b and Supplementary Fig. 3b). The field of view was positioned on the cross-section and points with high variance were obtained to tune the automatic focus/stigmatism functions (Fig.3c). In the last step, the acquisition started with a crisp focus (Supplementary Fig. 3c) and the system acquired section after section (slice and view). For 3D data acquisition the FIB was operated at 1.5 nA with the SEM and the FIB operating simultaneously ^55^. The images were acquired in analytical mode (1.5 kV, 700 pA) using the energy-selective back-scattered electron (EsB) detector at 1100 V grid voltage. The dwell time was set to 10 µs/pixel, with a line average of 1 and a 8 nm pixel size. The FOV was set to 25 *µm* x 15 *µm* (XY frontface) and images were collected for a ROI of 25 *µm* x 30 *µm* (XY, plane parallel to the surface) at 200 nm intervals for COP phenotypes, resulting in 8×8×200 nm voxels (XYZ)).

During the acquisition, an additional process was launched to monitor the status of the imaging (Supplementary Fig. 3b). This monitoring was in charge of automatically placing the region where the autofocus and autostigmatism (*AFAS*) routines had to be executed. The *AFAS* routines happened at periodic 45 minutes intervals and were executed in a region of the cell where high contrast could be found. This was achieved using a Harris corner detector ^29^ combined with clustering: the clustering with a higher number of corners is the candidate point for *AFAS*.

In addition, the imaged x-y region of interest was tracked in the y direction to prevent undesirable drifting. Cross-correlation was employed to find the relative difference between consecutive sections. The upper layer of the resin covered with gold was detected using changes in entropy and a drift >25% of the total y size, was corrected at 0.5 *µm* increments, maintaining the region of interest in the desired field of view. After the acquisition of one position was completed, the stage of the FIB-SEM was moved to the next ROI, and started a new acquisition.

#### Stereology

All stereological measurements were performed using IMOD^56^. From every siRNA treatment, a minimum of two cells typical for the individual treatment and very different from the negative control were selected by the image analysis pipeline. The FIB-SEM images were acquired throughout the cell of interest with a spacing of 200 nm and a random starting point producing 10-20 evenly spaced sections per cell. Golgi cisternae were defined as membranous structures devoid of ribosomes with a threefold length to breadth ratio. Golgi stack profiles were defined as any assembly of cisternal membranes and the total area enclosed by them ^57^. The volume of Golgi stack, (*V* (Golgi stack); Supplementary Fig. 4c) was estimated using a point counting-based Cavalieri estimator by applying a regular point lattice that yielded 100 to 200 points hits over Golgi stack profiles per typical section stack ^57^.

Our FIB-SEM sections were prepared orthogonal to monolayer substratum in a haphazard (random) orientation relative to the analyzed cell, and therefore comprise vertical sections on which the surface density of the Golgi cisternae in stack volume can be estimated by counting intersections (I) arrays of cycloid line probes ^38^. As cisternal membrane morphology was often indistinct, Golgi cisternae profiles were defined as a single line bisecting the cisternal membrane profile. Mean number of cisterna was estimated using the ratio of (intersections with Golgi cisterna; I(cist)) to (intersections with the Golgi stack profile face; I(stack face)).

