## Supplemental information for "CLEM*Site*, a software for automated phenotypic screens using light microscopy and FIB-SEM"

### Supplementary Figures

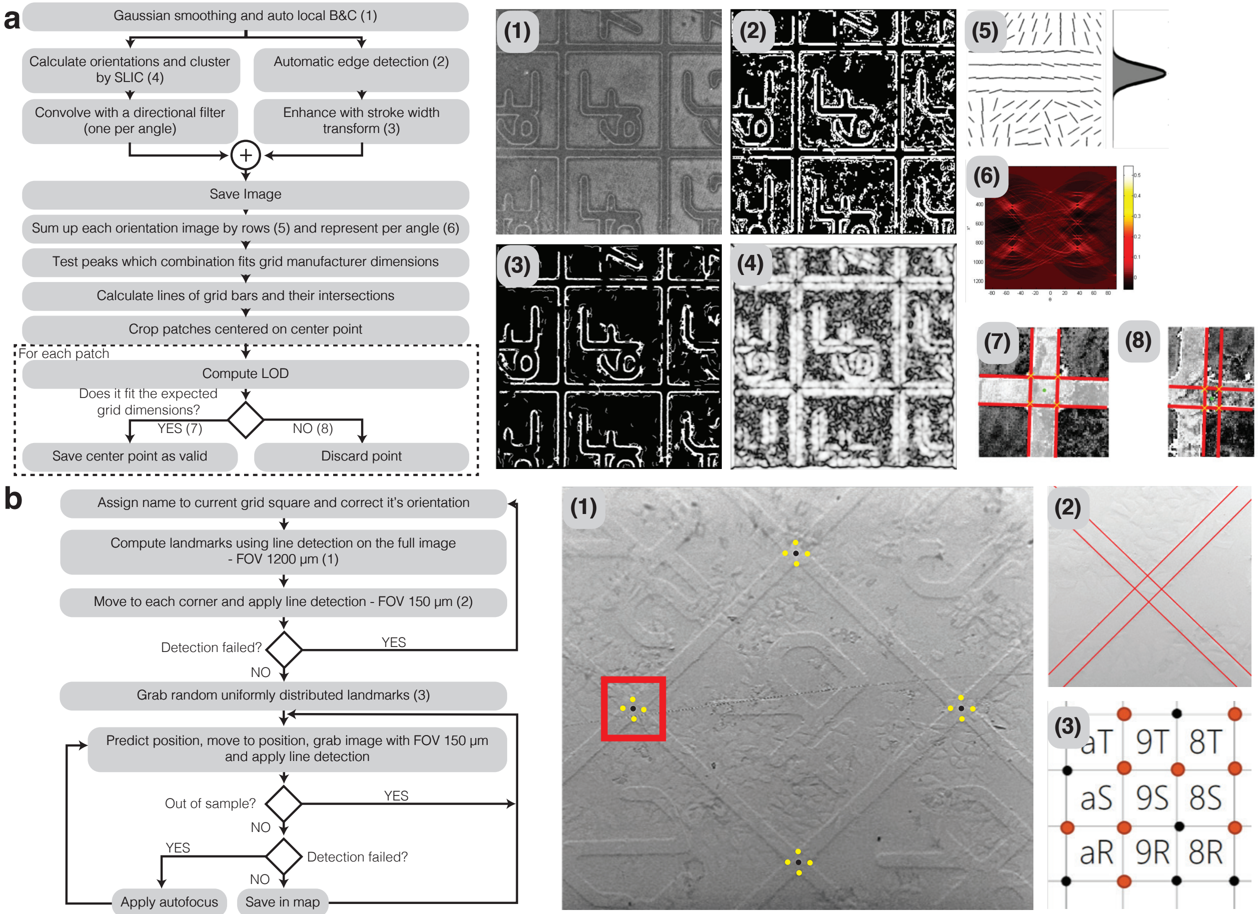


**Supplementary Figure 1:**

Line detection and landmark recognition. (a) Schematic of the line detection algorithm. Each step is illustrated with the corresponding output image: (1) Reflected light image of the coordinate system is smoothed and the brightness and contrast automatically balanced with adaptive histogram equalization. (2) Automatic edge detection is performed using Canny edge detection. (3) Image edges are enhanced with stroke width transform to facilitate the recognition of the alphanumerical pattern. (4) Pixel gradient orientations are extracted and homogenized in superpixels (SLIC). (5) The image resulting from 4 is convolved by every angle from 0 to 180 degrees, and all the rows summed up forming a projection. (6) From 5, peaks are found using non maxima suppression and tested to find the best fit to the grid dimensions according to the manufacturer. Each peak is a line detected in the image and can be plotted back in the original image. By calculating all the intersections between lines, the grid bar crossings can be found. (7) For each bar crossing, a refinement is applied. First an area is cropped, and the patch is analyzed again to enable a higher detection accuracy of the lines. When the distance between intersections is not fitting the expected grid pattern, the landmark is not accepted (8). (b) Schematic of the algorithm used by the *Navigator* module to find landmarks in the SEM, in order to build a map based on the grid. In the first step, the SEM is positioned at a random square in the MatTek grid, as shown in (1). The software detects the corners (black dots) by detecting the line intersections of the square edges (yellow points). Each corner is refined by applying the line detection (red lines) in a higher magnification view (2). In order to optimize the process, the detection procedure is applied to only a group of selected landmarks in the MatTek grid. By following a uniform random distribution (3), the same accuracy of the transformation can be achieved with less examined landmarks. If the line detection fails, an autofocus is applied once. If after a second round, the detection fails, the landmark position is flagged as blocked. In this way, landmark positions that fall outside the sample or are too damaged, are discarded from the final landmark map.

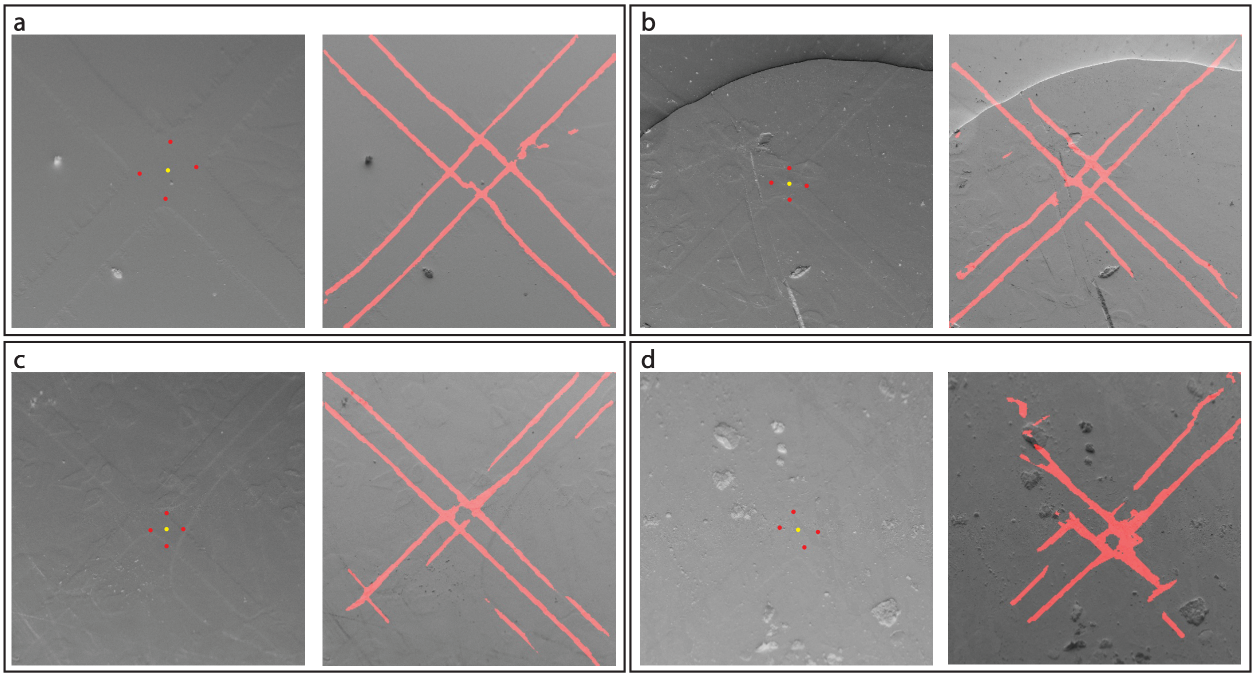


**Supplementary Figure 2:**

Cracks, scratches and dirt on the surface make the landmark detection difficult and more error prone. For each square, the left image shows the final detection, with the yellow dot representing the detected center position of the crossing and the red points the corners of the crossing. The right image is the same image (inverted), where the red pixels represent the probability of being a grid edge as detected by the neural network.(a) The sample is in perfect state, (b) a crack present in the upper part might affect the predicted accuracy of the overall map, even if the detection is good. (c) scratches can be the cause false positives for the grid detection. (d) dirt and other material residues, in this case from silver painting (usually used around the sample border to derive charges), might mislead the detection algorithm and increase the final error.

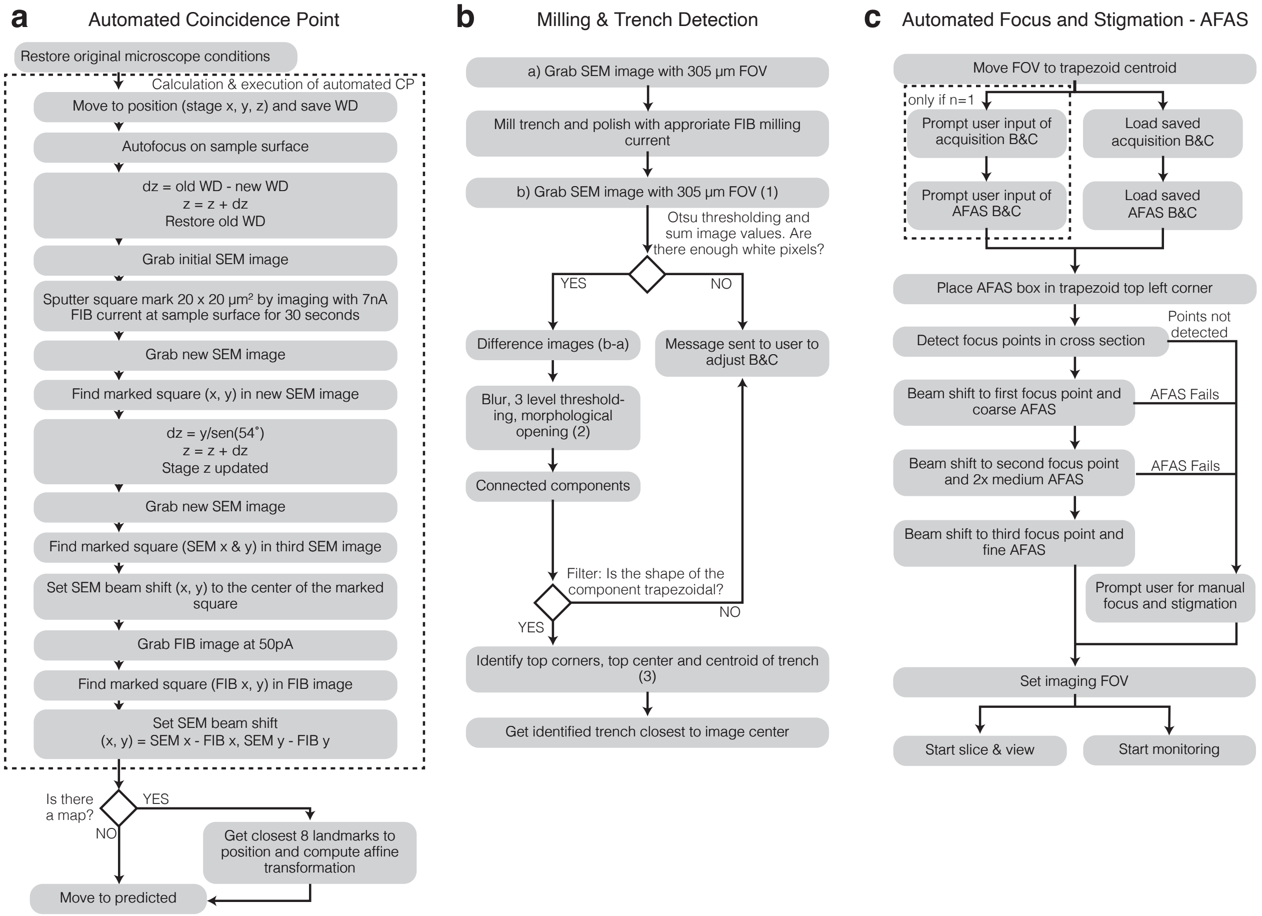


**Supplementary Figure 3: Automatic workflow setup for data acquisition in the FIB-SEM (*Multisite*).** (a) Detailed flowchart of the algorithm used for the Automated Coincidence Point. This procedure is executed before each target cell is acquired. The boxed part indicates the instructions belonging to the coincidence point (CP) calculation. Upon completion and to prevent error from any shift caused by the CP calculation, when a stored map of landmarks is present the closest 8 landmarks are used to compute a local transformation that will re-estimate the cell position. WD - Working distance, dz - difference in z position, SEM x, SEM y - stage position coordinates x and y using the SEM detector. FIB x, FIB y - stage position coordinates x and y using the FIB detector. In both cases, pixel coordinates from the image are translated to stage position coordinates given by the center position of the image. (b) Flowchart of the algorithm used for Milling & Trench Detection. Numbers (1), (2) and (3) correspond with images (1), (2), (3) on Figure 3b. (c) Flowchart of the routine used for Automated Focus and Stigmation (*AFAS*) prior to acquisition. In the automation routine, for the first cell acquired (n=1), the user must decide the brightness and contrast (B&C) of the sample. Values of B&C will be stored for future acquisitions. After choosing an optimal B&C, the goal is to start with a crisp image with a good focus and stigmation set of values. The core *AFAS* routine is provided by *ZEISS Atlas 5* software and is triggered in a reduced window from the full field of view (FOV) at different magnifications, from lower to higher. At each magnification, high complexity regions are found to be the center of the window where the *AFAS* is applied. If this routine fails to find a good focus before starting to acquire, which could happen in exceptionally damaged samples, the user is prompted to focus manually and the values of focus taken as reference for the next acquisition.

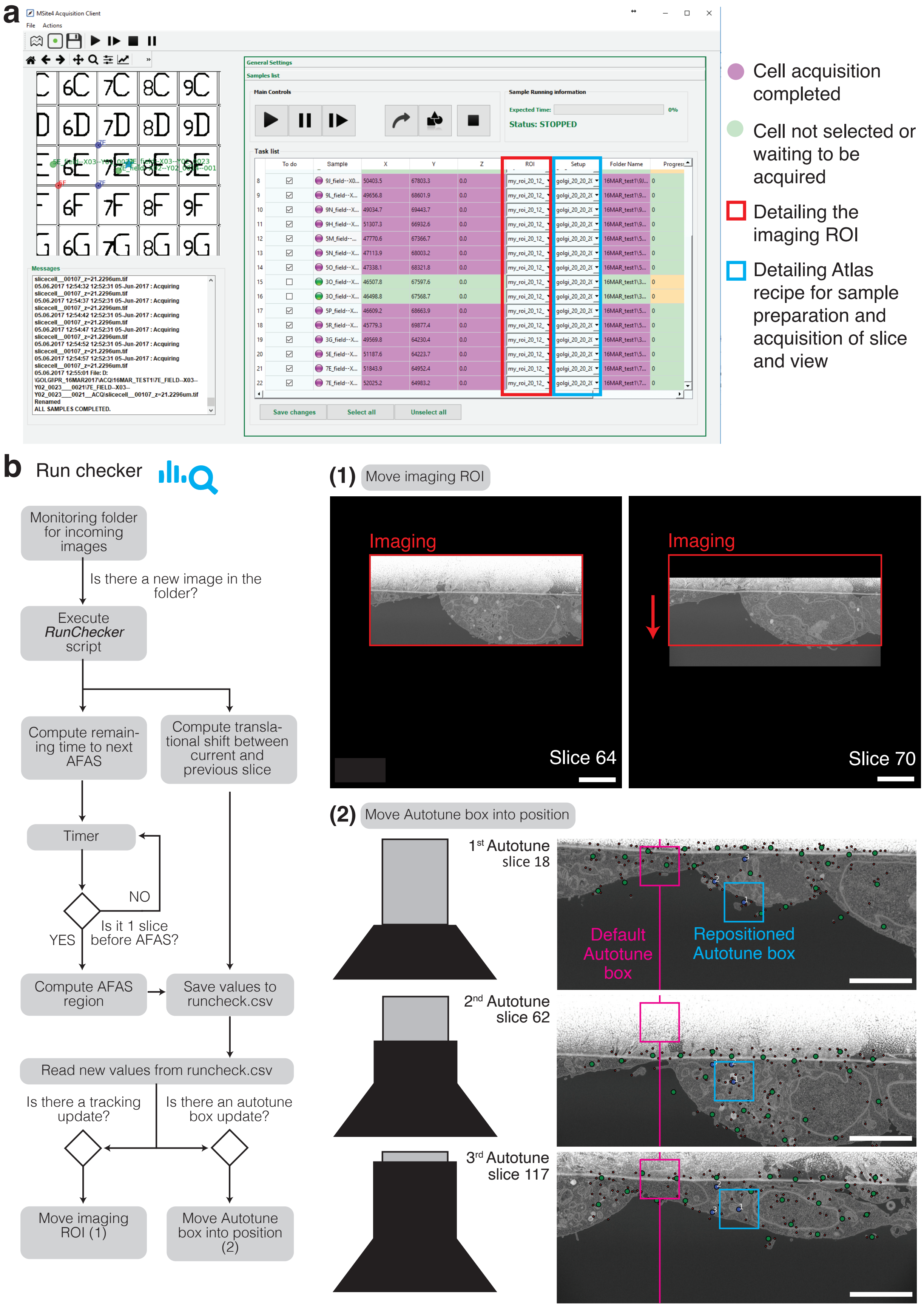
a
MSite4 Acquisition Client
General Settings
Samples list
Main Controls
Sample Running information
Status: STOPPED
Task list
Messages
ALL SAMPLES COMPLETED.
Save changes
Select all
Unselect all
Cell acquisition completed
Cell not selected or waiting to be acquired
Detailing the imaging ROI
Detailing Atlas recipe for sample preparation and acquisition of slice and view
b
Run checker
Monitoring folder for incoming images
Is there a new image in the folder?
Execute *RunChecker* script
Compute remaining time to next AFAS
Compute translational shift between current and previous slice
Timer
NO
YES
Is it 1 slice before AFAS?
Compute AFAS region
Save values to runcheck.csv
Read new values from runcheck.csv
Is there a tracking update?
Is there an autotune box update?
Move imaging ROI (1)
Move Autotune box into position (2)
(1) Move imaging ROI
Imaging
Slice 64
Imaging
Slice 70
(2) Move Autotune box into position
1st Autotune slice 18
Default Autotune box
Repositioned Autotune box
2nd Autotune slice 62
3rd Autotune slice 117

**Supplementary Figure 4: CLEM*Site*-EM interface and *Run Checker* details.** (a) Screenshot of the CLEM*Site*-EM interface to outline the details of the software User Interface (UI). In the top left panel, a map depicts targets (green) and landmarks (blue if SEM stage coordinates are matched with light microscopy stage coordinates, red if no match is present). Bottom left: a messaging console is used to display the communications with the server and which instructions are sent to the microscope. The right panel displays the list of all the targets to be acquired. The list presents which targets are already acquired (purple) and which ones are intact (green). Targets can be selected or deselected by ticking the *"To Do"* check box of the first column. In addition, for each target it is possible to decide the size of the section imaged from the total 3D volume milled by the FIB-SEM (ROI, red outline) and the *ZEISS Atlas 5* recipes for the actual acquisition (Setup, blue outline). (b) Flowchart of the logic applied by the *Run Checker* module. This module becomes active once a run starts. During the progression of the acquisition, the FOV carries a translational shift that has to be tracked and corrected continuously. In this module, the routine calculates the translation between two consecutive frames and makes a decision to move the imaging ROI if the sample has drifted (1). The same is applied to the position of the Autotune Box (small window where the *AFAS* is applied, magenta and blue squares) which is moved into a new position before a new *AFAS* is executed (2). In this case, the image is analyzed to find optimal positions for the Autotune Box.

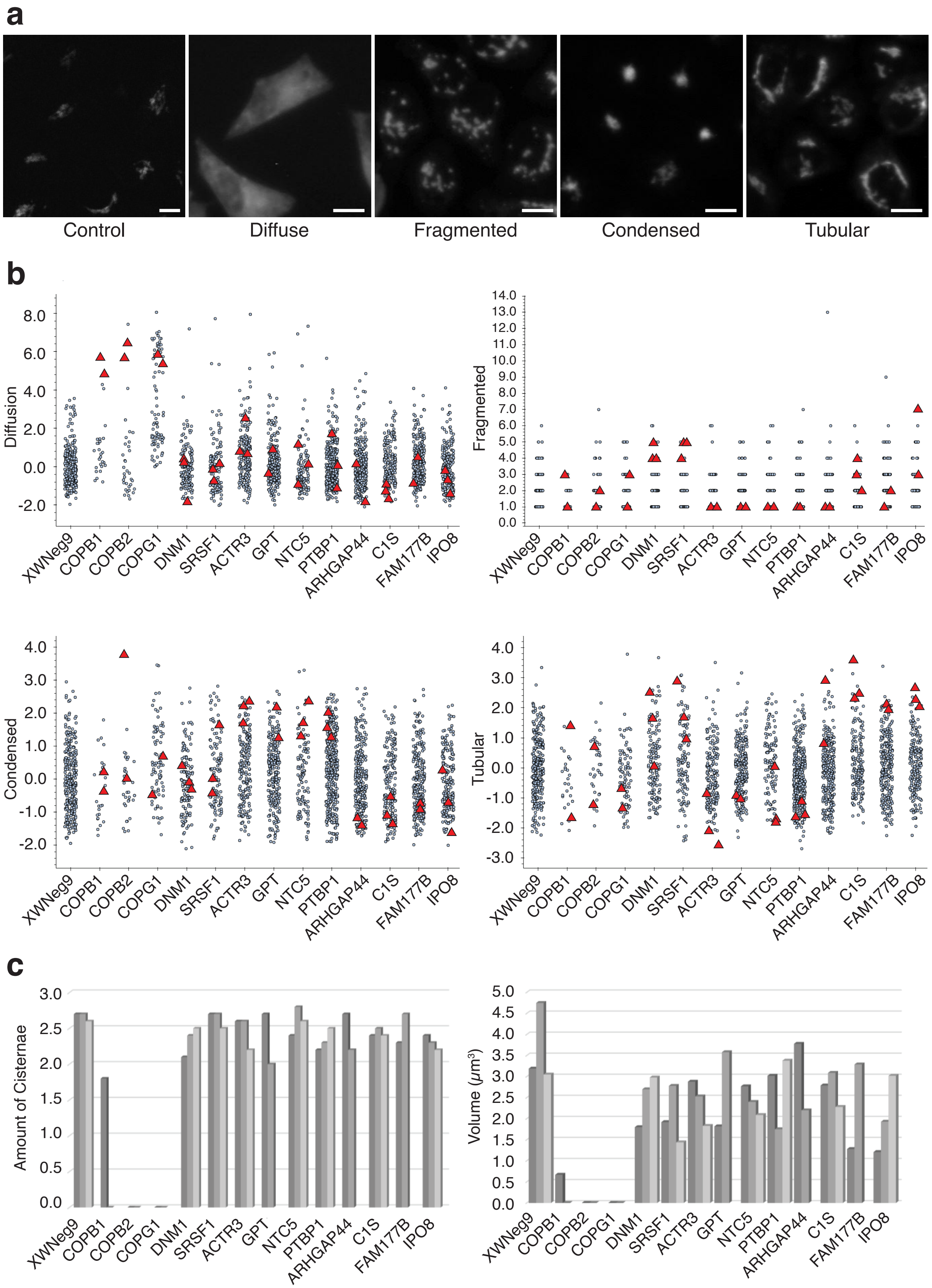
a
Control
Diffuse
Fragmented
Condensed
Tubular
b
Diffusion
Fragmented
Condensed
Tubular
XWNeg9
COPB1
COPB2
COPG1
DNM1
SRSF1
ACTR3
GPT
NTC5
PTBP1
ARHGAP44
C1S
FAM177B
IPO8
c
Amount of Cisternae
Volume (μm³)

**Supplementary Figure 5: Phenotype description and stereological quantification of chosen cells for the entire workflow.**

(a) Illustrations of the different Golgi phenotypes revealed by the GalNac-T2 GFP signal: control, diffuse (COPG1), fragmented (DNM1), condensed (ACTR3) and tubular (IPO8). Scale bars: control, 5 $\mu m$, rest, 10 $\mu m$. (b) Scatter plots of computed features measuring the strength for each phenotype. Each gray dot represents the feature value associated with one cell. In the x axis displays the corresponding siRNA treatment. Diffuseness, condensation and tubularity values are normalized with respect to the control (XWNeg9). Fragmentation illustrates the number of fragments detected in the Golgi apparatus. Red triangles highlight each one of the selected cells for the CLEM experiment (a total of 33). (c) Stereological quantification was applied on FIB-SEM images of the corresponding cells to measure number of cisternae (left) and the volume (right) of the Golgi apparatus. Each bar represents the value measured for one cell, grouped by siRNA treatment. Since the sample size is very small (n=2 or n=3 per treatment), the screen was oriented exclusively to find large effects. Knockdowns of the COP proteins (COPB1, COPB2, COPG1), revealed a disappearance of the Golgi stacks (thus, no cisternal volume can be measured) replaced by a large accumulation of small vesicles. No obvious morphological differences were found upon other siRNA treatments with respect to the control cells.

**Supplementary Table 1:**

Table 1: RMSE of targeting position

| | 1<br>RMSE Global | 2<br>RMS Local (estimated) | 3<br>RMS Local (measured) |
|---|---|---|---|
| S1 Spots (n=53) | 6.44 ± 4.3 | 4.53 ± 3.4 | 4 ± 1.9 |
| S2 Spots (n=46) | 9.62 ± 5.1 | 4.26 ± 3.1 | 4.94 ± 3.5 |
| S1 COPB1 (n=33) | 18.76 ± 11.5 | 12.73 ± 10 | 12 ± 4.3 |
| S2 COPB1 (n=47) | 20.56 ± 13.5 | 14.98 ± 13.5 | 9.86 ± 6.5 |
| Avg and std (n=179)* | 13.21 ±6.2 | 8.71± 5.2 | 7.7±4.4 |

* except for last column, in which n = 40
(1) RMSE of global transformation using all samples in an affine transform, (2) RMSE using a transformation involving only local samples (8 closest landmarks), (3) RMSE measured by manual registration of images. Only n=10 of the images were used per experiment

**Supplementary Table 2: Description of the 32 siRNA spots added to one dish in Case Study 1, referring to figure 4.**

| | Gene name | Full name | Gene description | siRNA ID (silencer select Ambion) | Sense siRNA Sequence | Antisense siRNA Sequence | Golgi phenotype |
|---|---|---|---|---|---|---|---|
| 1* | COPB2 | coatomer protein complex, subunit beta 2 | Coatomer complex required for budding from Golgi membranes and essential for the retrograde Golgi-to-ER transport of dilysine-tagged proteins | s17738 | CGAUGUAUCUCCUAGGCUAtt | UAGCCUAGGAGAUACAUCGtc | Diffuse |
| 2 | WDR75 | WD repeat domain 75 | Ribosome biogenesis factor | s38530 | CAGCUAGCAAAGAUGGUUAtt | UAACCAUCUUUGCUAGCUGta | Condensed[†] |
| 3* | DNM1 | dynamin 1 | Dynamin subfamily of GTP-binding proteins, involved in clathrin-mediated endocytosis and other vesicular trafficking processes | s144 | GCAGUUCGCCGUAGACUUUtt | AAAGUCUACGGCGAACUGCtg | Fragmented |
| 4* | COPG1 | coatomer protein complex, subunit gamma | Coatomer complex required for budding from Golgi membranes and essential for the retrograde Golgi-to-ER transport of dilysine-tagged proteins | s22431 | CGUCGGAUGUGCUACUUGAtt | UCAAGUAGCACAUCCGACGga | Diffuse |
| 5 | C1S | complement component 1, s subcomponent | Encodes a serine protease, which is a major constituent of the human complement subcomponent C1. | s2157 | CCAAGUCCCAUACAACAAAtt | UUUGUUGUAUGGGACUUGGaa | Tubular[†] |
| 6 | DENND4C | DENN/MADD domain containing 4C | Guanine nucleotide exchange factor (GEF) activating RAB10. Promotes the exchange of GDP to GTP. | s31214 | GUUUGGACCUUCCUAGUAAtt | UUACUAGGAAGGUCCAAACgt | Fragmented[†] |
| 7* | IPO8 | importin 8 | nuclear protein import | s20635 | CAUUCAACAUUCACGAAAAtt | UUUUCGUGAAUGUUGAAUGga | Tubular |
| 8 | SRSF1 | splicing factor, arginine/serine-rich 1 | Ensures the accuracy of splicing and regulating alternative splicing. | s12727 | GCAUCUACGUGGGUAACUUtt | AAGUUACCCACGUAGAUGCgg | Fragmented[†] |
| 9 | XWNeg9 | | negative control | s444246 | UACGACCGGUCUAUCGUAGtt | CUACGAUAGACCGGUCGUAtt | |
| 10 | NT5C | 5',3'-nucleotidase, cytosolic | Catalyzes the dephosphorylation of the 5' deoxyribonucleotides (dNTP) and 2'(3')-dNTP and ribonucleotides | s195191 | GCUUUUUCCUGGACCUGGAtt | UCCAGGUCCAGGAAAAAGCcc | Condensed[†] |
| 11* | ACTR3 | ARP3 actin-related protein 3 homolog | ATP-binding component of the Arp2/3 complex | s19642 | GGACGAGAUAUAACAUAUUtt | AAUAUGUUAUAUCUCGUCCtg | Condensed |

| | | | | | | | |
|---|---|---|---|---|---|---|---|
| 12 | PTBP1 | polypyrimidine tract binding protein 1 | Involved in mRNA metabolism and transport | s11436 | GCAUCACGCUCUCGAAGCAtt | UGCUUCGAGAGCGUGAUGCgg | Condensed† |
| 13* | DNM1 | dynamin 1 | Dynamin subfamily of GTP-binding proteins, involved in clathrin-mediated endocytosis and other vesicular trafficking processes | s144 | GCAGUUCGCCGUAGACUUUtt | AAAGUCUACGGCGAACUGCtg | Fragmented |
| 14 | FAM177B | family with sequence similarity 177, member B | | s53382 | AGACUACUCCUAAAAGGAUtt | AUCCUUUUAGGAGUAGUCUtt | Tubular† |
| 15 | PTBP1 | polypyrimidine tract binding protein 1 | Involved in mRNA metabolism and transport | s11436 | GCAUCACGCUCUCGAAGCAtt | UGCUUCGAGAGCGUGAUGCgg | Condensed† |
| 16 | ARHGAP44 | Rho GTPase activating protein 44 | GTPase-activating protein (GAP) that stimulates the GTPase activity of Rho-type GTPases. | s19218 | AGAACCUCUUAUGACCUUUtt | AAAGGUCAUAAGAGGUUCUgg | Tubular† |
| 17 | XWNeg9 | | Negative control | s444246 | UACGACCGGUCUAUCGUAGtt | CUACGAUAGACCGGUCGUAtt | |
| 18* | ACTR3 | ARP3 actin-related protein 3 homolog | ATP-binding component of the Arp2/3 complex | s19642 | GGACGAGAUAUAACAUAUUtt | AAUAUGUUAUAUCUCGUCCtg | Condensed |
| 19 | SRSF1 | splicing factor, arginine/serine-rich 1 | Ensures the accuracy of splicing and regulating alternative splicing | s12727 | GCAUCUACGUGGGUAACUUtt | AAGUUACCCACGUAGAUGCgg | Fragmented† |
| 20 | C1S | complement component 1, s subcomponent | Encodes a serine protease, which is a major constituent of the human complement subcomponent C1 | s2157 | CCAAGUCCCAUACAACAAAtt | UUUGUUGUAUGGGACUUGGaa | Tubular† |
| 21* | IPO8 | importin 8 | nuclear protein import | s20635 | CAUUCAACAUUCACGAAAAtt | UUUUCGUGAAUGUUGAAUGga | Tubular |
| 22 | WDR75 | WD repeat domain 75 | Ribosome biogenesis factor | s38530 | CAGCUAGCAAAGAUGGUUAtt | UAACCAUCUUUGCUAGCUGta | Condensed† |
| 23 | NT5C | 5',3'-nucleotidase, cytosolic | Catalyzes the dephosphorylation of the 5' deoxyribonucleotides (dNTP) and 2'(3')-dNTP and ribonucleotides | s195191 | GCUUUUUCCUGGACCUGGAtt | UCCAGGUCCAGGAAAAAGCcc | Condensed† |
| 24 | FAM177B | family with sequence similarity 177, member B | | s53382 | AGACUACUCCUAAAAGGAUtt | AUCCUUUUAGGAGUAGUCUtt | Tubular† |
| 25* | COPB1 | coatomer protein complex, subunit beta 1 | Coatomer complex required for budding from Golgi membranes and essential for the retrograde Golgi-to-ER transport of dilysine-tagged proteins | s3371 | GGUCUGUCAUGCUAAUCCAtt | UGGAUUAGCAUGACAGACCtt | Diffuse |

| | | | | | | | |
|---|---|---|---|---|---|---|---|
| 26 | ARHGAP44 | Rho GTPase activating protein 44 | GTPase-activating protein (GAP) that stimulates the GTPase activity of Rho-type GTPases. | s19218 | AGAACCUCUUAUGACCUUUtt | AAAGGUCAUAAGAGGUUCUgg | Tubular† |
| 27 | XWNeg9 | | negative control | | | | |
| 28 | GPT | glutamic-pyruvate transaminase (alanine aminotransferase) | Plays a key role in the intermediary metabolism of glucose and amino acids. | s6103 | CAGUUCCACUCAUUCAAGAtt | UCUUGAAUGAGUGGAACUGcg | Condensed† |
| 29 | AURKB | Aurora B Kinase | positive transfection control | s17612 | UCGUCAAGGUGGACCUAAAtt | UUUAGGUCCACCUUGACGAtg | Multinucleated |
| 30 | GPT | glutamic-pyruvate transaminase (alanine aminotransferase) | Plays a key role in the intermediary metabolism of glucose and amino acids. | s6103 | CAGUUCCACUCAUUCAAGAtt | UCUUGAAUGAGUGGAACUGcg | Condensed† |
| 31 | DENND4C | DENN/MADD domain containing 4C | Guanine nucleotide exchange factor (GEF) activating RAB10. Promotes the exchange of GDP to GTP. | s31214 | GUUUGGACCUUCCUAGUAAtt | UUACUAGGAAGGUCCAAACgt | Fragmented† |
| 32 | KIF11 | kinesin family protein 11 | positive transfection control | s7903 | GACUGAUCUUCUAAGUUCAtt | UGAACUUAGAAGAUCAGUCtt | No cells |

*Genes with highlighted examples in Figure 4

† Genes represented a variety of listed phenotypes giving a trend for this phenotype
